# The human hepatocyte TXG-MAPr: WGCNA transcriptomic modules to support mechanism-based risk assessment

**DOI:** 10.1101/2021.05.17.444463

**Authors:** Giulia Callegaro, Steven J. Kunnen, Panuwat Trairatphisan, Solène Grosdidier, Marije Niemeijer, Wouter den Hollander, Emre Guney, Janet Piñero Gonzalez, Laura Furlong, Yue W. Webster, Julio Saez-Rodriguez, Jeffrey J. Sutherland, Jennifer Mollon, James L. Stevens, Bob van de Water

## Abstract

Mechanism-based risk assessment is urged to advance and fully permeate into current safety assessment practices, possibly at early phases of drug safety testing. Toxicogenomics is a promising source of comprehensive and mechanisms-revealing data, but analysis tools to interpret mechanisms of toxicity and specific for the testing systems (e.g. hepatocytes) are lacking. In this study we present the TXG-MAPr webtool (available at https://txg-mapr.eu/WGCNA_PHH/TGGATEs_PHH/), an R-Shiny-based implementation of weighted gene co-expression networks (WGCNA) obtained from the Primary Human Hepatocytes (PHH) TG-GATEs dataset. Gene co-expression networks (modules) were annotated with functional information (pathway enrichment, transcription factor) to reveal their mechanistic interpretation. Several well-known stress response pathways were captured in the modules, are perturbed by specific stressors and show preserved in rat systems (rat primary hepatocytes and rat *in vivo* liver), highlighting stress responses that translate across species/testing systems. The TXG-MAPr tool was successfully applied to investigate the mechanism of toxicity of TG-GATEs compounds and using external datasets obtained from different hepatocyte cells and microarray platforms. Additionally, we suggest that module responses can be calculated from targeted RNA-seq data therefore imputing biological responses from a limited gene. By analyzing 50 different PHH donors’ responses to a common stressor, tunicamycin, we were able to suggest modules associated with donor’s traits, e.g. pre-existing disease state, therefore connected to donors’ variability. In conclusion, we demonstrated that gene co-expression analysis coupled to an interactive visualization environment, the TXG-MAPr, is a promising approach to achieve mechanistic relevant, cross-species and cross-platform evaluation of toxicogenomic data.

## Introduction

Mechanism-based risk assessment is the gold standard for safety assessment of drugs and chemicals, but is resource- and time-intensive (Lanzoni et al., 2019). Establishing mechanisms generally requires extensive experimentation, leveraged literature data collected over years, and is often based on preclinical (animal) studies, which may or may not translate to human (Bailey & Balls, 2019; Clark & Steger-Hartmann, 2018; World Health Organization, 2017). Approaches that rapidly reveal mechanistic detail of preserved responses between animal and human would advance next generation risk assessment and impact both predicting and monitoring toxicity (Rivetti et al., 2020). For example, early in development, drugs that cause preclinical toxicity are often discarded based on preclinical studies without knowing if a toxicity mechanism will translate to human, thus valuable compounds may be discarded. In addition, screening for safer compounds is hampered if testing assays do not interrogate compounds against a mechanism-related event and results are not interpreted in the context of organ-specific toxicity.

Liver is a primary target organ for drug and chemical toxicants due to its role in metabolism and disposition (J. Zhang & Venkat, 2020). Drug-induced liver injury (DILI) is a major cause of clinical liver failure (Björnsson, 2019; Reuben et al., 2016), drug attrition and black box warnings (Onakpoya et al., 2016; Solotke et al., 2018; Watkins, 2011). DILI manifests as a variety of clinical pathologies and may depend on genetic and environmental factors making its prediction a challenge for the current preclinical testing paradigm (Koido et al., 2020). Likewise, hepatotoxicity and hepatocarcinogenesis are major concerns for environmental exposures (Colombo et al., 2019; Yorita Christensen et al., 2013). Both non-genotoxic and genotoxic carcinogens often produce hepatoxicity prior to the emergence of liver tumors in longer term preclinical studies (Karin & Dhar, 2016). In both situations, hepatotoxicity can be regarded as a multistep, multicellular disease process, where an initial molecular stress is followed by a series of cellular key events that couple the initial stress to an apical endpoint observable as a pathology, e.g. hepatotoxicity or liver tumors. However, in the absence of a mechanism that links cellular (stress) events to a pathology, risk assessment is typically based on the apical endpoint which can take months or years to develop.

Toxicogenomics, the study of transcriptome responses in toxicology, is a promising tool for a comprehensive analysis of toxicity to enable mechanism-based risk assessment (Liu et al., 2019). Current toxicogenomic approaches mainly rely on the analysis of Differentially Expressed Genes (DEGs) or enrichment analysis using annotated pathway analysis tools (Barel & Herwig, 2018). Such an approach depends on ontologies with a high degree of redundancy and capture current knowledge of toxicology and general biology. These approaches are useful but biased toward genes that are well annotated and are not designed to interpret mechanisms of toxicity applied to specific testing systems (e.g. hepatocytes). For these and other reasons, and despite several decades of research, toxicogenomics has not produced the anticipated transformation of current safety assessment standard practice (Vahle et al., 2018).

It is known that groups of genes expressed downstream of a (stress-responsive) transcription factor will show co-expression (Yin et al., 2021). For this reason, gene co-expression analysis has been applied to toxicogenomic datasets for rat liver (Podtelezhnikov et al., 2020; J. J. Sutherland et al., 2018), but not for human hepatocytes up to now. These co-expressed gene sets mediate the cellular response to stress, providing mechanistic information on the cellular processes and key events involved in adaptation and progression. Herein, we used weighted gene co-expression network analysis (WGCNA) to identify sets of co-expressed genes (termed ‘modules’) using the large TG-GATEs toxicogenomic dataset for primary human hepatocytes (PHH) as a surrogate to address hepatotoxicity and gene expression in the human context (Igarashi et al., 2015; B. Zhang & Horvath, 2005). Gene modules were deployed in an R-Shiny analysis framework accessible via an interactive website, the PHH TXG-MAPr (https://txg-mapr.eu/WGCNA_PHH/TGGATEs_PHH/) to facilitate rapid visualization and mechanistic interpretation of transcriptomic data. Gene module were annotated with external annotation resources (e.g. pathway enrichment and transcription factor modulation), which are useful in identifying known stress response pathways and modules populated by novel genes responsive to specific stressors. Using endoplasmic reticulum (ER) stress as a case study, we show that canonical ER stress response genes are captured in gene co-expression modules, including novel guilt-by-association genes. We identified ER stress as an early event in cyclosporine A (CSA)-induced toxicity and showed that ER stress modules respond similarly between CSA and the prototypical ER stressor, tunicamycin. ER stress and other known stress response pathways which were captured in the PHH modules, are conserved across species and test systems. Stress response modules correlate in clusters with defined biological functions triggered by compound exposure. We showed that datasets of different sources can be analyzed with the PHH TXG-MAPr tool for mechanistic interpretation. Finally, we leveraged our PHH WGCNA modules to evaluate donor-to-donor variability upon tunicamycin exposure and identified biological sources of variations. Our results show that co-expression analysis of toxicogenomic data using the PHH TXG-MAPr framework can support ready and human-relevant next generation mechanism-based risk assessment.

## Materials and Methods

### Gene expression data processing

#### TG-GATEs data

Microarray data from primary human hepatocytes (PHH) in the TG-GATEs repository were downloaded and jointly normalized using the Robust Multi-array Average (RMA) method within the *affy* R package. To map probe sets to Entrez IDs, the BrainArray chip description file (CDF) version 20 was used (http://brainarray.mbni.med.umich.edu/Brainarray/Database/CustomCDF/genomic_curated_CDF.asp, HGU133Plus2 array version). Under this annotation, every gene is defined by a single probe set. This resulted in 19363 unique probe sets, each mapped to a single gene. The TG-GATES (TG) repository contains 941 PHH treatments, where each treatment is defined as a combination of compound, time and concentration. Samples treated with vehicles are available for each individual time point. For each experimental condition (compound/timepoint combination), the *limma* R package was used to calculate log2 fold-change values, which was performed by building a linear model fit and computing the log-odds of differential expression by empirical Bayes moderation. TG-GATEs rat primary hepatocytes (RPH) and rat *in vivo* liver datasets have been analyzed following the same steps using suitable BrainArray CDFs (version 19, Rat2302 array version). Additional human hepatocytes datasets for uploading into the TXG-MAPr tool were first downloaded from Gene Expression Omnibus (GEO: https://www.ncbi.nlm.nih.gov/geo/): GSE83958; GSE45635; GSE53216; GSE74000; GSE13430; GSE104601. The first four datasets were processed similarly to the TG-GATEs PHH dataset, because the samples were analyzed with the same microarray platform (Affymetrix HGU-133 Plus 2). The GSE13430 and GSE104601 datasets were generated from the Agilent Human 1A Microarray (V2) G4110B and SurePrint G3 Human GE v2 8×60K microarray platforms, respectively. These datasets were analyzed with GEO2R using the provided NCBI gene annotation. In the case of duplicated measurements (probes) for a single gene, the most significant probe (adjusted p value) was used for uploading the data.

#### TempO-Seq S1500+ set PHH data

The inter-individual variability in chemical-induced ER stress signaling and module activation was evaluated by using available TempO-Seq data of PHHs derived from 50 individuals which were exposed to a wide concentration range of tunicamycin (Niemeijer et al., in press, Mav et al., 2020). In short, plateable cryopreserved PHHs derived from 50 individuals (KaLy-Cell, Plobsheim, France) were plated in 96 wells BioCoat Collagen I Cellware plates (Corning, Wiesbaden, Germany) at a density of 70,000 viable cells per well. Cells were allowed to attach for 24 hours and treated with tunicamycin (Sigma) at a concentration range of 0.0001 to 10 µM for 8 or 24 hours. After exposure, cells were lysed with 1x TempO-Seq lysis buffer (BioSpyder) and stored at −80°C. Lysates were analyzed at BioSpyder (Carlsbad, CA, USA) using the TempO-Seq technology in combination with the S1500+ gene set (Mav et al. 2018). Data analysis is reported in Niemeijer et al. 2021, here briefly summarized: experiments having library size of raw counts lower than 100,000 counts were filtered out. Raw counts were normalized with DESeq2 normalization (Love et al., 2014), log2 transformed and analyzed with BMD express (Phillips et al., 2019).

### Weighted gene co-expression analysis

In order to identify co-expressed genes from the PHH data, we used the *WGCNA* R package (Peter Langfelder et al., 2020) and applied it to a matrix consisting of 941 rows (PHH experiments) and 17500 columns (log2 fold change values for genes). Genes were included into the analyses when they showed FDR (BH) < 0.001 in at least one experiment. We created unsigned gene modules (i.e. grouping together co-induced and co-repressed genes), and selected the optimal soft power threshold maximizing both the scale-free network topology using standard power-law plotting tool in WGCNA (Langfelder & Horvath, 2008) and the exclusion of genes having low gene intensity in the DMSO controls (determined via t-tests). We selected 5 as the optimal soft-power parameter. By further refining modules built using WGCNA function to merge similar modules (those having correlation of their eigengene values ≥ 0.8), we obtained 398 modules containing 10,275 genes. As described in (Jeffrey J. Sutherland et al., 2016), for each experiment we calculated the eigengene score (EGs, or module score) which summarizes log2 fold change of their constituent genes. Briefly, this protocol consisted of performing PCA on the gene matrix of each module, normalizing the log2FC across the entire dataset using Z-score conversion. To simplify comparison between modules, the raw module score was normalized to unit variance (fraction between each module score and its standard deviation across the entire dataset) facilitating comparison across modules and across treatment. Therefore, the modules score indicates the level of activation or repression induced by a given treatment, when considering such changes in the context of the large collection of drug perturbations. Represented as Z-score, an eigengene score greater than +2.0 or smaller than −2.0 can be considered as a large (and relevant) perturbation in the context of the 941 experiments Although the EGs represent the overall activation or repression of the module gene members, each gene is ordered based on intra-modular connectivity with other genes. We calculated then the correlation between a module eigengene versus underlying genes’ log2 fold change across the 941 experiments (termed ‘corEG’). The gene with the highest correlation between log2FC and module EGs (so called ‘hub gene’) is the most representative of the entire module matrix and show stronger connection to the other module genes. Both positive and negative correlation with the EGs, corEG, were allowed, thus genes that are inversely correlated can be members of the same module. A negative corEG indicates an inverse relationship between the gene log2FCs and its module EGs. The matrix containing EGs across all treatments was organized into a folded hierarchical tree (dendrogram), based on Ward’s hierarchical clustering of pair-wise Pearson correlations for each module across all treatment conditions. Module locations on dendrogram branches were identified with a hierarchical anti-clockwise nomenclature system (Supplementary Figure S1A). Some module branches were elongated to separate module clusters and improve visualization (Supplementary Figure S1B). Compound correlation was calculated with Pearson and Spearman correlation using the module EGs across the entire set of treatment conditions (resulting in 941 instances); similarly, module correlation was calculated across all modules (resulting in 398 instances). Preservation between the modules structure obtained with the PHH TG-GATEs data set and the RPH TG-GATEs and rat *in vivo* liver data sets has been performed as follows: 1) rat gene IDs have been converted to human gene IDs with the Rat Genome Database (Smith et al., 2020), 2) preservation statistics have been calculated with the *WGCNA* R package and thresholds for interpretation were adopted by relevant literature (Langfelder et al., 2011). Briefly, a module showing Z summary >=2 is considered moderately preserved, >=10 highly preserved. A lower median rank indicates higher preservation.

### Pathway mapping and enrichment analyses

Over Representation Analysis (ORA) was performed via ConsensusPathDB (CPDB version 34), including the following databases: BioCarta, EHMN, HumanCyc, INOH, KEGG, NetPath, Reactome, Signalink, SMPDB, Wikipathways, UniProt, InterPro (Kamburov et al., 2013). GO enrichment was obtained with the R package topGO, algorithm = “classic”, statistic = “fisher” (Alexa & Rahnenführer, 2007). For both resources, we included in the tool enriched terms satisfying hypergeometric test p-value < 0.01. However, for module interpretation (Table 1 and Table S8) only the top 10 associated terms per modules were included, that corresponded to terms having always adjusted p value (FDR) for CPDB lower than 0.05. String DB (Szklarczyk et al., 2015) was used to obtain protein-protein interaction networks of specific modules. Edge weights are proportional to the combined score of the nodes that the edges connect (Szklarczyk et al., 2015). Graphical rendering was obtained with Cytoscape (Shannon et al., 2003) using the StringApp (http://apps.cytoscape.org/apps/stringapp) bridge component.

**Table 1.**
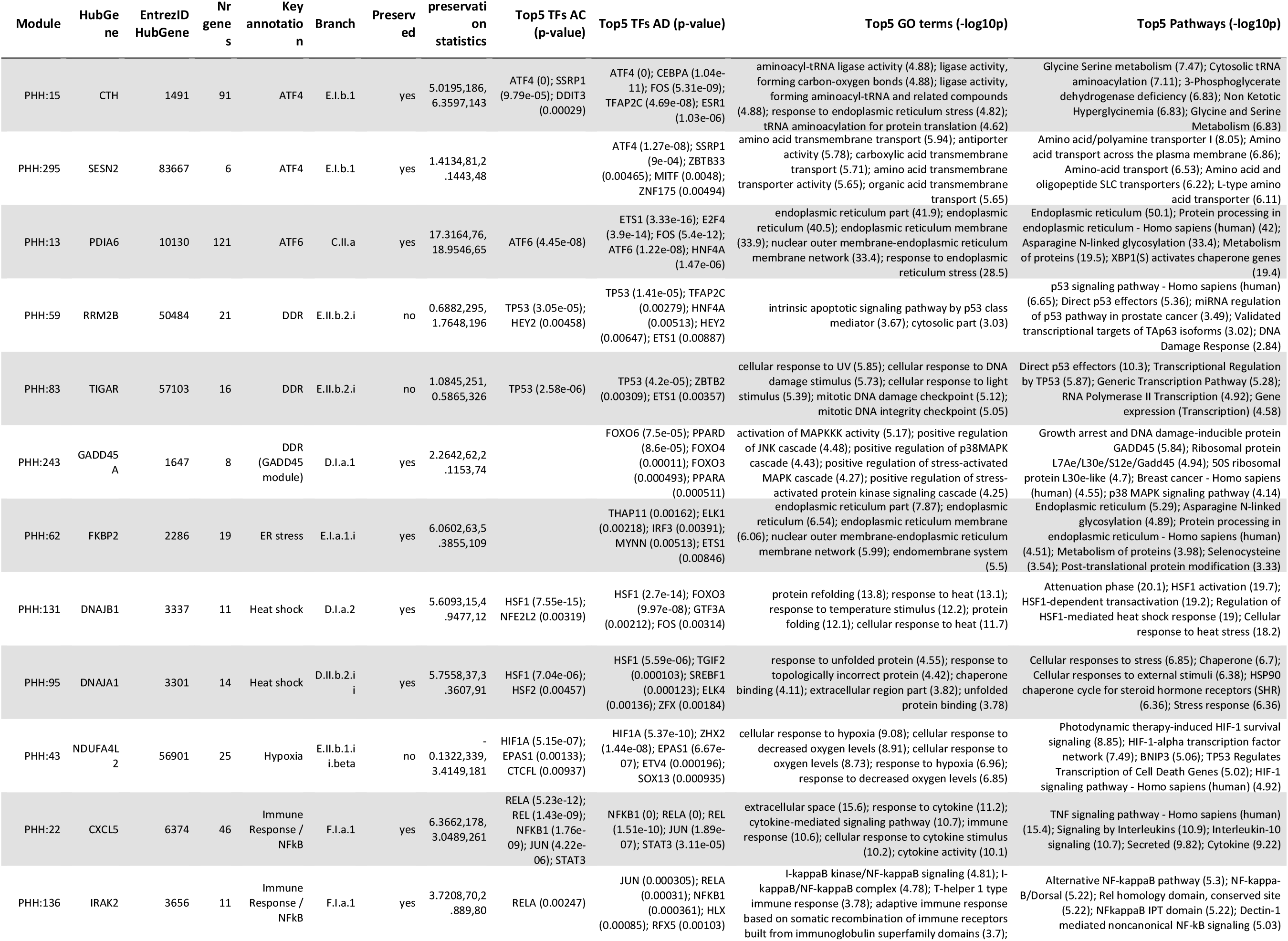
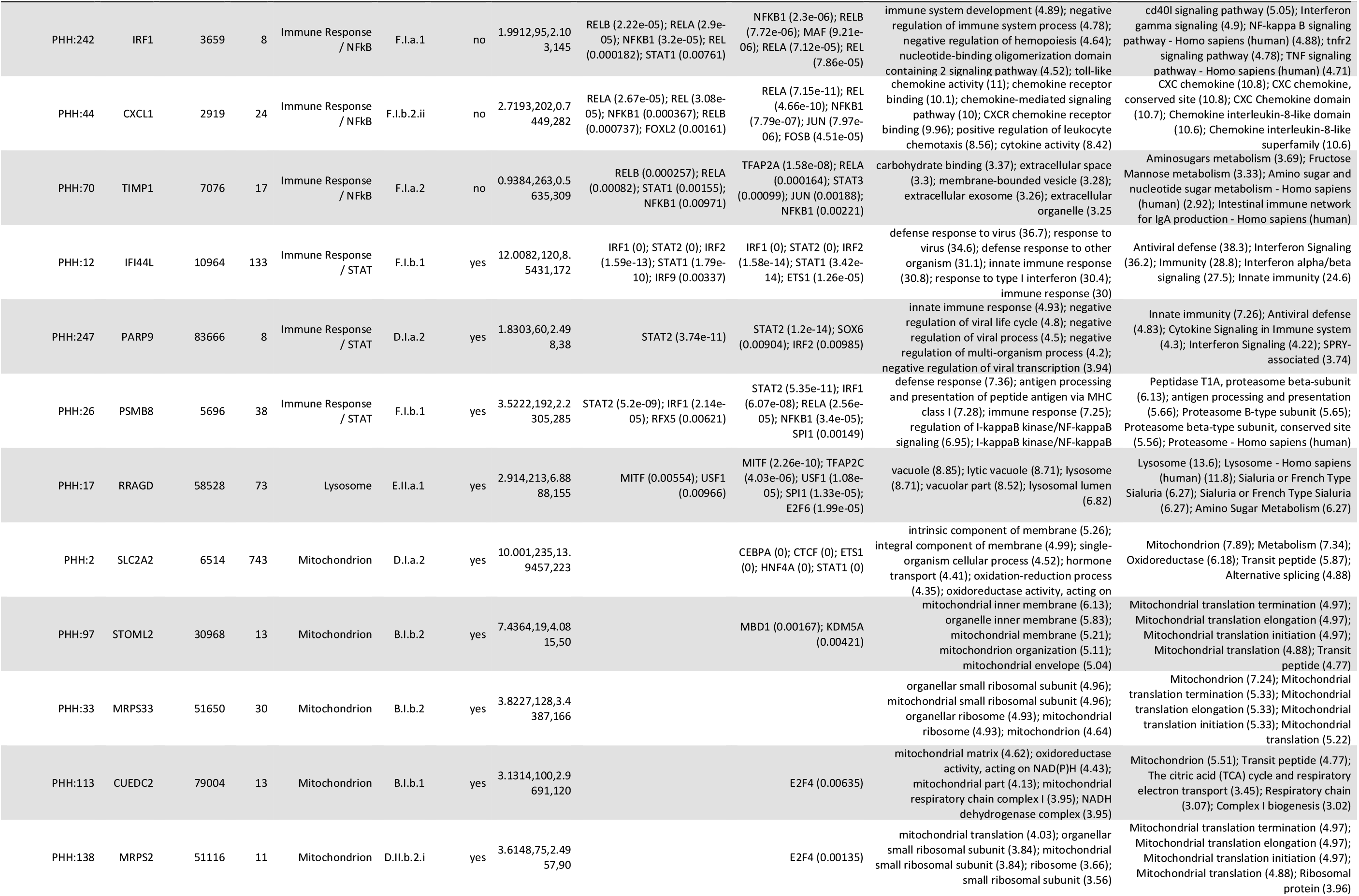
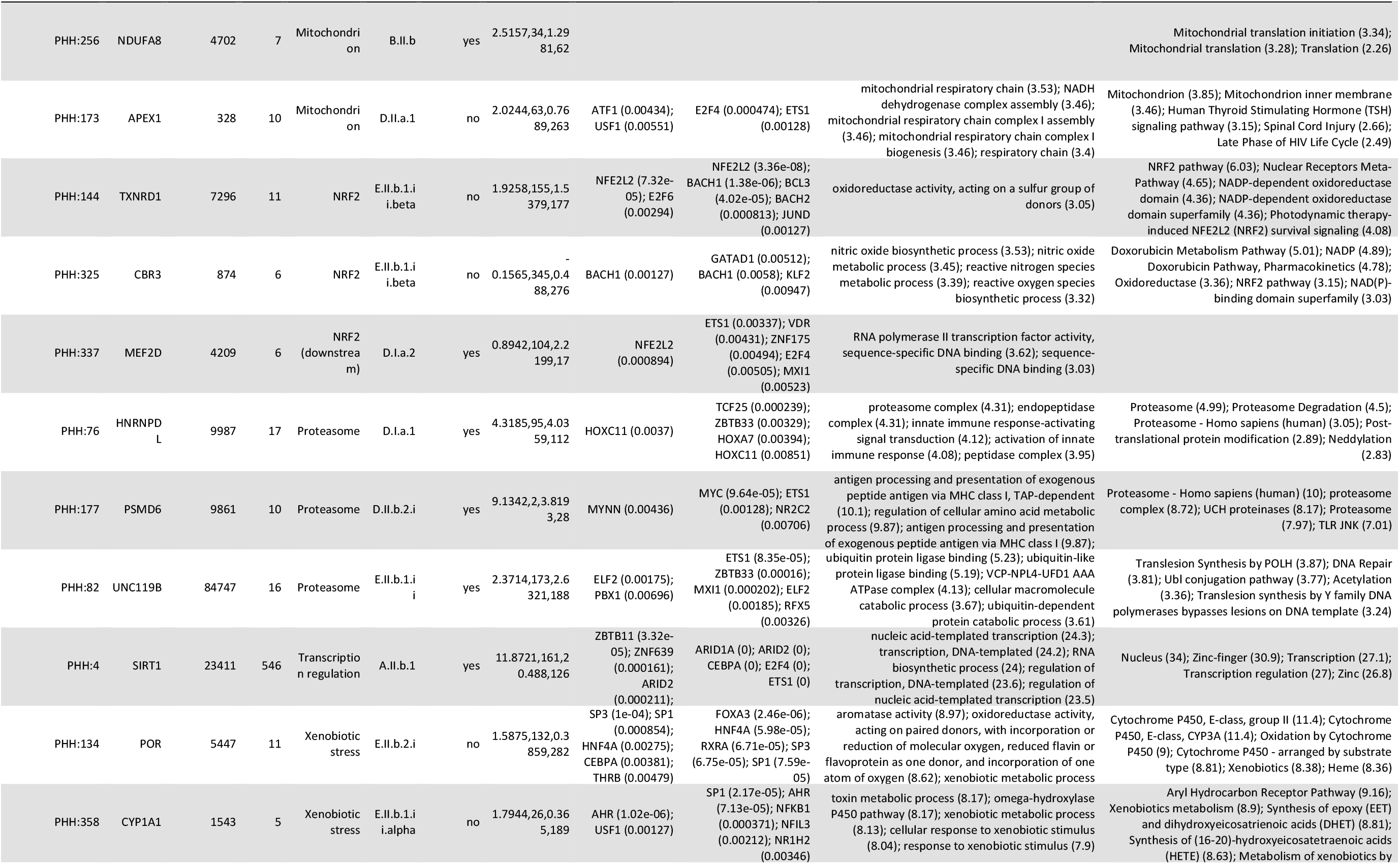
Overview of the properties of the modules associated with the stress response. Preservation statistics are, in order: Z summary PHHtoLIVER, MedianRank PHHtoLIVER; Zsum PHHtoRPH; MedianRank PHHtoRPH. More details can be found in Supplementary Tables S1-7.

### Transcription factor (TF) enrichment and TF activity scoring

A hypergeometric test was performed on gene members in each WGCNA module to identify its regulatory TFs using the function *phyper* within the *stat* package in R. The gene set of TFs and their regulated genes (regulons) are derived from DoRothEA (Garcia-Alonso et al., 2019) with two sets of confidence levels: the “high confidence” level comprises categories A, B and C while the “high coverage” level comprises categories A, B, C and D. The enriched TFs with p-value less than 0.01 were included in the study. In parallel, TFs’ activities were estimated as normalized enrichment scores using the function *viper* from the *viper* package (Alvarez et al., 2016) with two confidence sets of TF-regulon from DoRothEA as described. All parameter settings were assigned as in the original DoRothEA study (Garcia-Alonso et al., 2019).

### Web application TXG-MAPr

The user interface application of the TXG-MAPr tool has been implemented using the R-shiny package (RStudio Inc., 2014). The graphical part of the application has been implemented through the functionalities of the *ggplot2, ape, igraph, hclust* and *pheatmap* R-packages (Csardi & Nepusz, 2006; Kolde & Kolde, 2015; Paradis & Schliep, 2019; Wickham H, 2008). The credential system to log into the PHH TXG-MAPr tool is established using the *shinyauthr* package (https://github.com/PaulC91/shinyauthr). The PHH TXG-MAPr tool is available at https://txg-mapr.eu/WGCNA_PHH/TGGATEs_PHH/.

### Uploading new data files into TXG-MAPr

In order to calculate new EGs for each module from external data, first a modified Z-score is calculated by dividing the gene log2FC by the standard deviation of the gene log2FC across all TG-GATEs conditions, and is further weighted by the correlation eigengene score (corEG). (Eq. 1). New EGs for each module are calculated by summing the Z-scored gene log2FC values of the module genes (n = number of genes in a module), normalized by the standard deviation of the raw module score in the TG-GATEs dataset (Eq. 2). When data of certain genes in a module is not available in the uploaded dataset, then the Z-score for that gene is assumed to be zero, which may create an underestimation of the final module EGs.

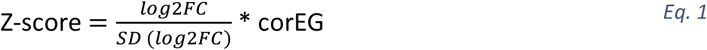

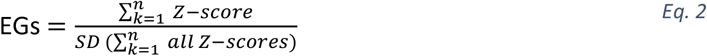

The new module EGs will be overlaid onto the PHH TXG-MAPr dendrogram and will be fully integrated into the web application for that particular session. Data will be removed when the session is closed.

### Data analyses

#### Cluster correlation of the expanded seed module set

Figures were made using R and the packages *ggplot2, pheatmap, igraph*. Heatmap in Figure 3 is obtained with the following steps: 1) selection of ‘seed modules’ i.e., modules having clear interpretation based on term enrichments and TFs associations (see Table 1). Because TG-GATEs data is a sparse matrix, i.e. not all the compounds have been tested at all concentration levels and time points, we removed the 2-hour experiments (missing for roughly 50% of the compounds) and applied imputation of the remaining missing value with Singular Value decomposition via the R package *pcaMethods* using the *svdImpute* function (Stacklies et al., 2007; Troyanskaya et al., 2001). 2) Pearson correlations between seed modules and all other preserved modules were calculated considering all remaining EGs from TG-GATEs (119 modules x 779 experiments). Modules were added in (termed ‘add-in modules’) if the (seed x target) correlation values were > 0.7 to yield the ‘expanded set’ of 87 modules. Hierarchical clustering was obtained by applying the Ward D2 method, using 1-cor as distance. In particular, we applied the WardD2 method, including the full implementation of the Ward clustering criterion. Note that the results of Ward’s agglomerative clustering are likely to delineate clusters that visually correspond to regions of high densities of points in PCA ordination, e.g. produce ‘round’ clusters (Murtagh & Legendre, 2014).

#### Targeted genes set analysis (S1500+)

To evaluate the quality of the genes belonging to the BioSpyder TempO-Seq S1500+ set, we tested for difference of mean absolute corEG with a random draw of genes from the complete pool of genes available in the TG-GATEs dataset. The randomly selected gene set have the same size as the overlap of the S1500+ set with the TG-GATEs gene set (n = 1830) and this process was repeated 10,000 times (Wilcoxon test), adjusting the p value with Bonferroni method. The heatmap in Figure 5 was obtained with the “*complete*” clustering algorithm applied to the Euclidean distance between samples. Donors module clusters were obtained with the same method as for TG-based module clusters (see paragraph above). The overlap between TG module clusters and donor module clusters was calculating using the *WGCNA* R package (Peter Langfelder et al., 2020) using the *overlap* function, which calculates the numerical overlaps between groups and quantify the significance by applying Fisher exact test. Donor traits (phenotypes) associations with modules responses after treatment of donor hepatocyte cultures were determined as previously described in (J. J. Sutherland et al., 2018). In brief: 1) different concentrations were analyzed separately, and only after 24 hours of exposure (generally higher transcriptomic changes). 2) for cluster to traits associations, cluster behaviors were calculated as mean of the EGs scores of the modules members of each cluster. Since only modules with high correlation populate a cluster, we expect that averaging their EGs would lead to a concordant score. To understand driver-modules for each association, we calculated module-to-trait relationships. 3) The association between cluster (or module) and the occurrence of a donor trait was quantified using Cohen’s d, a measure of effect size. Since the average absolute module eigengene (avgAbsEG), a measure of overall transcriptional activity, can have an effect size for traits associations, logistic regression was performed to determine the contribution of a given module in explaining the residual odds of toxicity after accounting for avgAbsEG, and the significance of the module represented as adjusted p-values (p-adj).

## Results

### The PHH TXG-MAPr: a tool to visualize and explore biological interpretation in primary human hepatocyte transcriptomic data

We applied WGCNA to the TG-GATEs primary human hepatocytes (PHH) dataset (Igarashi et al., 2015), which includes exposure data for 158 compounds at various times and concentration levels to identify the predominant networks of co-expressed genes (Figure 1A exemplifies the process for a subset of co-expressed genes). We obtained 398 networks (modules) of highly correlated genes. Modules are labelled with a number inversely proportional to the number of member genes; larger modules have smaller number labels. An eigengene score (EGs) was computed for each module as the first principal component of each module data matrix and further scaled (see Material and Methods). Thus, the EGs represents the trend (induction or repression) of the entire module based on the included genes. By performing WGCNA, the final data matrix was reduced to 941 columns (treatments) and 398 rows (module EGs) which corresponds to a 97.7% reduction in dimensionality of gene expression.

**Figure 1.**
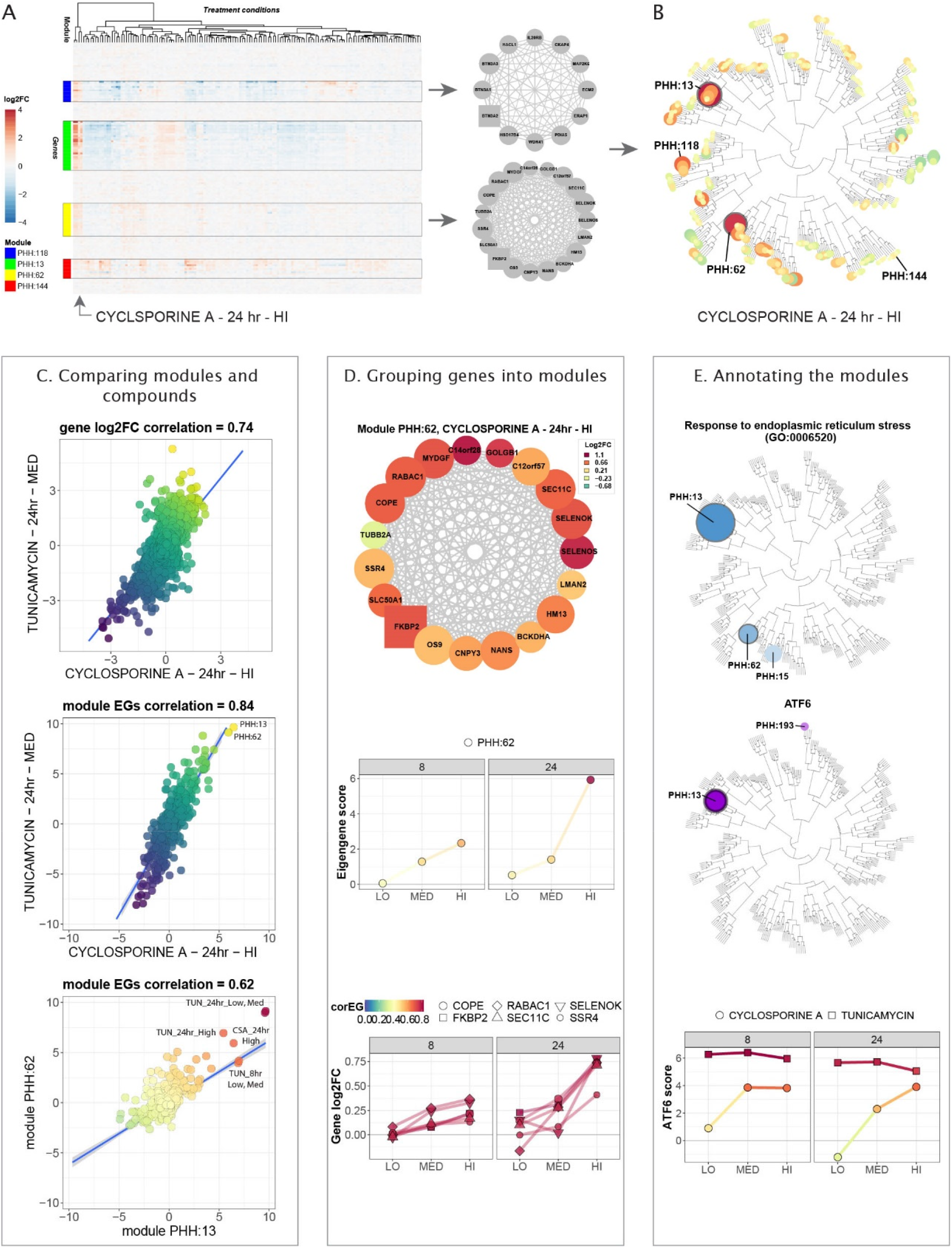
The PHH TXG-MAPr: an innovative tool to visualize and understand PHH toxicogenomic data. A) Example gene expression data matrix. Log2FC values of genes (rows) are shown for multiple treatment conditions (columns). Four groups of co-expressed genes (modules PHH:13, 62, 118 and 144), highlighted with boxes and color, show consistent patterns across the experimental conditions and are exemplified with the gene networks on the right side. Treatments are clustered using Euclidean distance to group conditions that regulate the same groups of co-expressed genes. B) Phylogenic tree view of module scores at 24hrs cyclosporine A exposure at HI dose level (6 µM). Highlighted modules are PHH:13, PHH:62 and PHH:118 which are strongly induced by CSA (orange/red circles), while module PHH:144 is unchanged (yellow circles). C) Comparing modules and compounds. Top: gene log2 fold change (log2FC) for cyclosporine A (HI dose = 6 µM, 24 hr) are plotted against gene log2FC for tunicamycin (MED dose = 2 µg/mL, 24 hr), showing a Pearson correlation of 0.74. Center: module EGs for cyclosporine A are plotted against module EGs for tunicamycin, showing a Pearson correlation of 0.84. Bottom: EGs for module PHH:62 are plotted against EGs for module PHH:13 (all compounds, concentrations and time points) and show a Pearson correlation of 0.62. Straight blue line represents a fitted linear model. Grey shades represent confidence interval. D) Grouping genes into modules. Top: gene network for module 62. Colors qualitatively represent log2FC upon treatment with cyclosporine A at HI dose (6 µM) for 24 hrs. Edges thickness is proportional to the adjacency value among the genes. The squared gene is the hub-genes. Center: module EGs profile for module PHH:62 at different cyclosporine A concentrations and time points (LO = 0.24 µM, MED = 1.2 µM, HI = 6 µM). Bottom: gene log2FC profile for hub-like genes belonging to module 62 at different cyclosporine A concentrations and time points. Color code of points and edges represents the correlation eigengene (corEG). E) Annotating the modules. Top: Phylogenic tree view with module color and size proportional to the p-value for the enrichment for the GO category response to endoplasmic reticulum stress (GO:0006520). Significantly enriched modules with p-values (<0.01) are displayed and modules PHH:13, 15 and 62 are highlighted in blue. Center: Phylogenic tree view with module color and size proportional to the p-value for the enrichment for the transcription factor ATF6. Significantly enriched modules for ATF6 with p-values (<0.01) are displayed and modules PHH:13 and 193 are highlighted in purple. Bottom: ATF6 activation score considering all its targets for cyclosporine A (LO = 0.24 µM, MED = 1.2 µM, HI = 6 µM) and tunicamycin (LO = 0.4 µg/mL, MED = 2 µg/mL, HI = 10 µg/mL).

To facilitate the analysis of transcriptomic information from module responses, we developed a module-based R-Shiny visualization and analysis framework, the PHH toxicogenomic (TXG) MAPr (PHH TXG-MAPr). The EGs-treatment data matrix was organized into a folded hierarchical tree, or dendrogram, based on Ward’s hierarchical clustering of pair-wise Pearson correlations for each module across all treatment conditions (Figure 1B, Supplemental Figure S1A-B). The dendrogram allows one to visually appreciate the induction or repression of modules EGs (red to blue color scale, respectively) and the proximity to modules with similar behavior (Pearson R; see Supplementary Figure S2 for dose- and time-response dendrograms for example compounds). We also incorporated module enrichment measures for biological pathways and ontologies, transcription factor-target pairs, compound similarity analysis and a variety of other useful functions for mechanistic analysis and data mining.

To illustrate the utility of the PHH TXG-MAPr analysis framework, we investigated the response of cyclosporine A (CSA), which was transcriptionally active in PHH and has both mild hepatotoxic and more severe renal toxicity potential (Rezzani, 2004). Strong induction of modules PHH:13 and PHH:62 can be seen 24 hours after treatment with 6 µM CSA (Figure 1B). Using the compound correlation functionality, we sampled all pair-wise Pearson correlation for all 398 module EGs across all treatment conditions; CSA shows highest similarity to tunicamycin (Pearson R = 0.84), suggesting a common mode of action (Figure 1C, center). Modules at the extremities of the correlation plot (PHH:13 and PHH:62 in the example) can be highlighted and tabulated to facilitate further analysis. Correlation between gene log2FC from the same treatment conditions had lower similarity (Pearson R = 0.74; Figure 1C, top). On average, across all TG-GATEs compounds the EGs-based max Pearson correlations were higher than max correlation based on gene log2FC (Supplementary Figure S1C), suggesting that analyzing gene networks (modules) improves the robustness of compound comparison of transcriptomic data by averaging out the variations of individual gene perturbations. Although PHH:13 and PHH:62 are in different branches they did show a Pearson correlation of 0.62 for all TG-GATEs treatment conditions (Figure 1C, bottom).

Tunicamycin (TUN) is a prototypical inducer of ER stress while there are literature reporting that CSA also perturbs ER functions (Foufelle & Fromenty, 2016; Van Summeren et al., 2013; Vickers et al., 2017). Therefore, we investigated modules PHH:13 and PHH:62 for genes associated with ER and ER stress. Module PHH:13 contains 121 genes, including well-known ER stress genes, like HSP90B1 (Grp94), SEL1L, PDIA6, HSPA5 (BiP/Grp78) and its co-chaperone DNAJB9 (Suppl. Table S2). Since PHH:13 was too large to visualize in a gene interaction plot, Figure 1D (top) shows the module plot for the smaller module PHH:62 (19 genes). This module contains genes involved in endoplasmic reticulum-associated degradation (ERAD), such as SELENOS and SELENOK, but also other ER resident genes with less clear connection to ER stress yet, such as FKBP2, SEC11C and SSR4 (Figure 1D, top). There is a clear time- and dose-dependent induction of module PHH:62 by CSA, which follows a similar trend for the log2FC induction of the hub-like (highest corEG) module genes (Figure 1D, center and bottom).

Using the PHH TXG-MAPr module enrichment functionalities, we investigated the enrichment of biological processes and pathways (see Methods) as well as transcription factors (TFs) enrichment using the DoRothEA gene set resource (Garcia-Alonso et al., 2018). Results were overlaid on the module dendrogram (Figure 1E). Not surprisingly, modules PHH:13 and 62 showed the lowest p-values for GO-CC term “*endoplasmic reticulum*” as well as terms related to endoplasmic reticulum (ER) stress amongst others (Figure 1E, top). Module PHH:13 was also enriched for transcription factor ATF6 target genes (Figure 1E, center) consistent with the presence of more highly annotated ER stress genes. To determine if PHH:13, the primary module enriched for ATF6 gene targets, reflected general ATF6 activation, we calculated the ATF6 activation scores for all ATF6 target genes in the 10275 TXG-MAPr genes (see more details on DoRothEA in the Materials and Methods section) and we observed that ATF6 scores also showed time- and dose-dependent activation for both CSA and TUN (Figure 1E, bottom).

Using the PHH TXG-MAPr analysis framework, we were able to rapidly identify the activation of an ATF6 regulated ER stress response as an early event following cyclosporine A exposure. Data underpinning these functionalities, and others not noted here, are accessible in a tabular format in the supplementary materials (Supplementary Tables S1-S7). The dedicated PHH TXG-MAPr application is available at https://txg-mapr.eu/WGCNA_PHH/TGGATEs_PHH/.

### Module preservation in the PHH-TXG-MAPr

Preservation statistics can be used to determine if networks’ node-edge relationships defined in one biological system are preserved in another (Langfelder et al., 2011). We evaluated network preservation of PHH modules versus rat primary hepatocyte (RPH) and *in vivo* rat liver TG-GATEs datasets using two different preservation statistics: Z-summary and Median Rank (see Methods, Figure 2A-B, Supplementary Table S8, Supplementary Figure S3). Z-summary preservation statistic shows higher dependency on modules size, as noted in literature; large modules such as PHH:13 are highly preserved (Figure 2A, color scale) (Langfelder et al., 2011). The complementary Median Rank statistic (Figure 2B) is less dependent on module size and can assign high preservation ranking (quantified with low numerical scores) to small modules. Therefore, we considered preserved modules as those with Z-summary preservation score for both systems (RPH and rat liver) higher than 2 or ranked in the top 100 modules for the Median Rank statistics for both systems (for the latter, arbitrary threshold corresponding to 25% of all the PHH modules). This selection led to a set of 102 (26%) preserved modules from the PHH TXG-MAPr (Supplementary table S8), which include the UPR modules PHH:13 and PHH:62.

**Figure 2.**
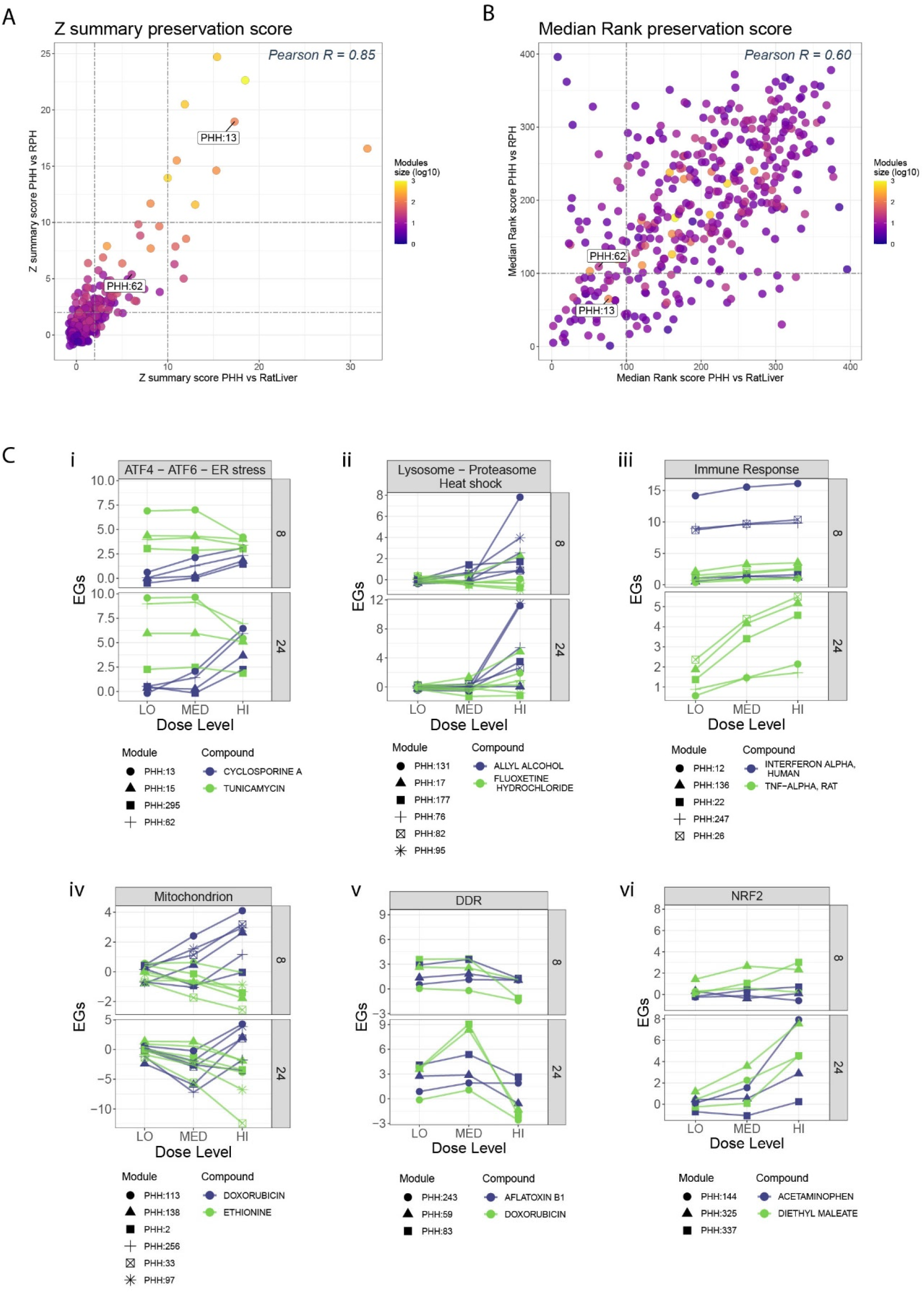
Some PHH WGCNA modules are preserved in rat systems and connect to stress response pathways. A) Z summary preservation score plot. Z summary preservation values of PHH modules in Rat in vivo Liver data (x-axis) are plotted against Z summary preservation values of PHH modules in Rat Primary Hepatocytes data (y-axis). Modules are colored based on their size (log10 transformed) and modules PHH:13 and PHH:62 are labeled. Higher scores imply better preservations. B) Median Rank preservation score plot. Median Rank preservation values of PHH modules in Rat in vivo Liver data (x-axis) are plotted against Median Rank preservation values of PHH modules in Rat Primary Hepatocytes data (y-axis). Modules are colored based on their size (log10 transformed) and modules PHH:13 and PHH:62 are labeled. Lower scores imply a higher rank and greater preservation. C) Dose- and time-response EGs plots of modules representing stress response pathways. Modules are grouped by stress processes (roman letters) and their responses upon treatment with representative compounds is shown in dose-response (x axis), faceted by time. Modules are represented by different symbols, while compounds by different colors.

### Stress response pathways represented by PHH modules

Pathway information facilitates mechanism-based risk assessment (Krewski et al., 2020). Therefore, we used the TXG-MAPr enrichment functionalities to identify modules representing cellular stress response pathways typically involved in toxicity and activated by stress-responsive TFs. These include the ER stress response and the larger integrated stress response (ISR), as well as inflammatory pathway, mitochondrial response, oxidative stress and the DNA damage responses, all of which were previously shown to be relevant for identifying the modes of action of compound-induced toxicity (Wink et al., 2014). We focused on modules which 1) were identified by consistent enriched terms and TF associations with p-values < 0.01 and FDR < 0.05, and 2) show a response to prototypical compounds (Figure 2C, Table 1). Below are the selected stress responses pathways represented by PHH modules which we aim to highlight in this study:

#### ER stress and ISR

As noted above, modules PHH:13 and PHH:62 are enriched for terms associated to endoplasmic reticulum localization, UPR and ERAD as well as transcriptional regulation by ATF6, all of which are components of the larger ISR (Hetz et al., 2020; Ron & Walter, 2007). We also looked for modules regulated by ATF4, a transcription factor downstream of the ISR hub, and IF2α, a regulator of translation after stresses including ER stress, starvation, and viral infection (Pakos-Zebrucka et al., 2016). Modules PHH:15 and PHH:295 were enriched for ATF4 gene targets, and terms associated with amino acid metabolism and transport, respectively (Table 1), which are processes that are regulated by ATF4 (Hetz et al., 2020). All four modules respond to cyclosporine A and tunicamycin (Figure 2C, panel i).

#### Heat-shock, proteasome and lysosome

In response to proteotoxic stress, damaged proteins are bound to members of heat shock-inducible chaperone system (heat shock response), which facilitate removal of damaged proteins by lysosomes and/or proteasome degradation. Modules PHH:177, PHH:76 and PHH:82 (proteasome) and PHH:131, PHH:95 (heat shock) are preserved and respond to treatment with allyl alcohol (Mandrekar et al., 2008) (Figure 2C, panel ii, Table 1). Module PHH:17 (annotated for GO:CC lysosome and includes SQSTM1) is also found to be preserved and is activated by fluoxetine, a known phospholipidosis inducer (Breiden & Sandhoff, 2019) (Figure 2C, panel ii, Table 1).

#### Immune response

Immune response pathways including inflammatory mediators are activated in hepatocytes during liver injury and in disease states (Campos et al., 2020; Woolbright, 2017). Immune response and inflammation terms are enriched in several preserved modules (PHH:12, PHH:247, PHH26, PHH:22 and PHH:136) that respond to inflammatory agents such as interferon-α and TNF-α (Figure 2C, panel iii, Table 1). Modules PHH:242, PHH:44 and PHH:70 are also annotated for immune response but not preserved (Table 1). Compound regulation and module enrichment terms identify subgroups of immune response modules. STAT signaling modules (PHH:12, PHH:247) are strongly induced by interferon, while NF-kB signaling modules (PHH:22, PHH:136) are mainly induced by TNFα. Module PHH:26 is induced by both stimuli, but also has a mixed NF-kB and STAT annotation.

#### Mitochondria

Mitochondria are an important target for hepatotoxic chemicals and mitochondrial injury is commonly used to screen for hepatotoxic potential (Rana et al., 2019; Weaver et al., 2020; Yang et al., 2015). Mitochondrial damage is commonly assessed by changes in respiration and the mitochondrial membrane potential. Less is known about the regulation of genes coding for mitochondrial proteins in response to toxic stress. Several preserved PHH modules are found to be annotated with mitochondria-related and mitochondria-component specific terms (PHH:113, PHH:138, PHH:2, PHH:256, PHH:33, PHH:97). These modules respond positively to doxorubicin (Osataphan et al., 2020) and are found to be repressed by ethionine (L. Zhang et al., 2020) (Figure 2C, panel iv, Table 1). Notably, PHH:33 and PHH:97 were annotated by terms relating to regulation of mitochondrial ribosome translational control and mitochondrial organization. Module PHH:173 shows enrichment for mitochondria genes, but is not preserved (Table 1).

#### DNA-damage & oxidative stress

Modules PHH:59 and PHH:83 are enriched for TP53-regulated genes and annotated for terms consistent with the DNA damage response (DDR). The NFE2L2 (NRF2) associated modules PHH:144 and PHH:325 include key genes such as SRXN1 and NQO1 which are direct targets of NRF2, and enrich for terms such as oxidoreductase and glutathione metabolism. In contrast to the other preserved modules from Table 1, modules associated with NRF2 and TP53 activation are not well-preserved, but these modules do respond as expected to specific inducers consistent with interpretation as oxidative stress and DNA damage modules, respectively (Figure 2C, panels v and vi and Table 1). Only modules PHH:243 (includes GADD45A/B) and PHH:337 (associated with NFE2L2) are found to be preserved (Table 1).

The examples described above exemplify the utility of PHH TXG-MAPr for identifying modules relevant to known stress pathway useful for mechanistic interpretation and benchmarking preservation of co-expression across species and liver *in vivo*. Other cellular processes and pathways not discussed can be identified using the main enrichment terms and TFs associations of the whole preserved module set (Table S8).

### PHH stress response modules seed clusters of modules in an interaction map to describe mechanisms of toxicity

In toxicity pathways, early events are coupled to a cascade of perturbations that lead to an adverse outcome, e.g. cell death. Using the stress pathway modules as seeds, we created an interaction map (cluster correlation) between stress pathways and other highly preserved modules that cluster important cellular processes as well as preserved modules without clear annotation and identify off-diagonal interactions between important biological themes.

We used the well-annotated modules identified above (Table 1 and Table S8) as seeds, and identified preserved ‘add-in’ modules (with or without clear enrichment) if they had a Pearson correlation >0.7 with a seed module, calculated across all TG-GATEs PHH compound treatments. The final expanded set of 87 seed and add-in modules is in Table S8, identified by the column “Cluster”. Figure 3 shows a cluster correlation analysis of the expanded seed set module EGs based on all TG-GATEs compounds, together with representative compound responses. Eight distinct clusters emerged with modules annotated for similar processes forming macro groups; examples are described below. Note that within each group, modules with limited to no annotation have been included, suggesting they belong to the same response group.

**Figure 3.**
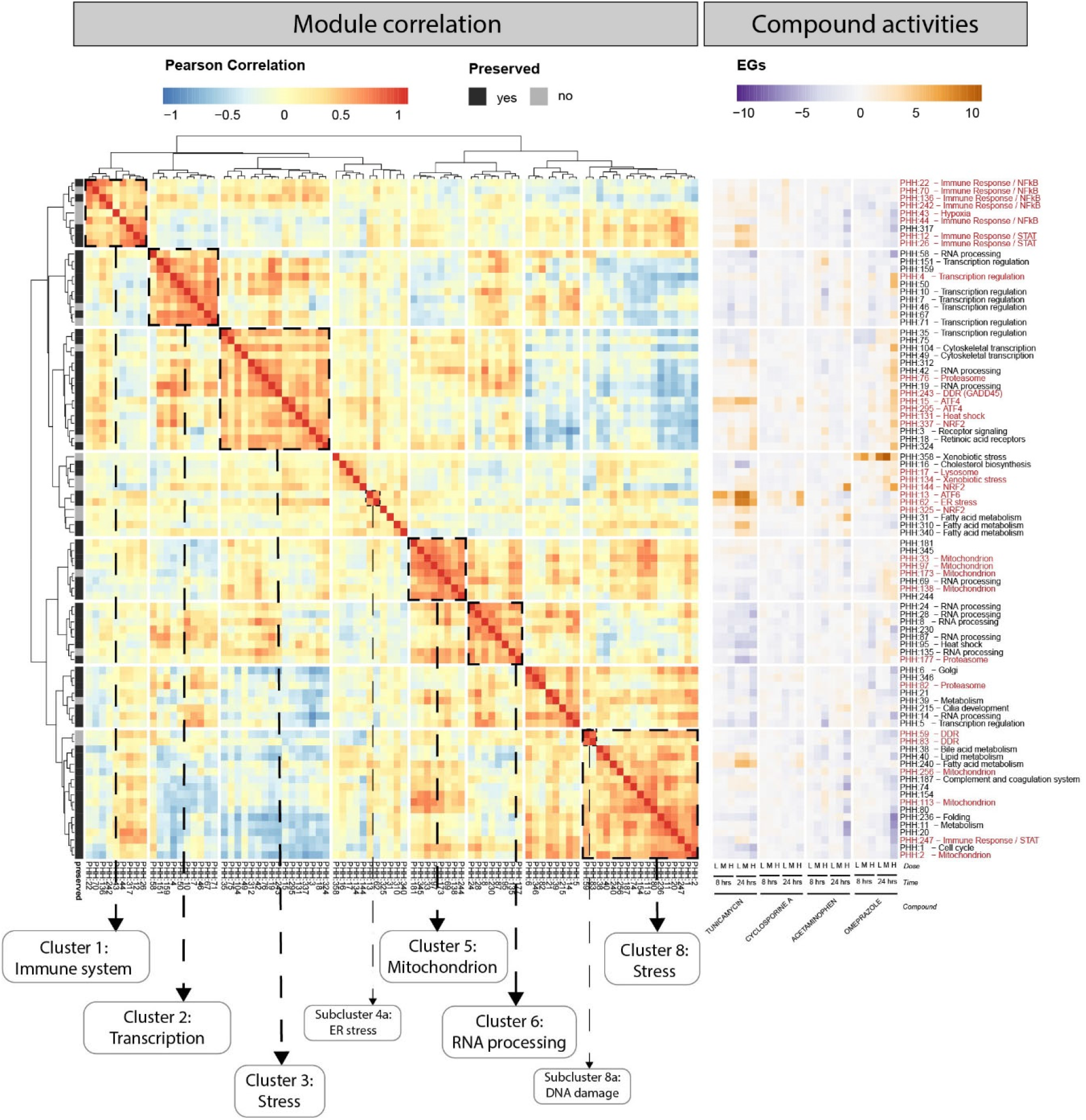
PHH stress modules interaction map. Cluster correlation matrix of the 87 modules correlating with well-annotated modules. On the left, modules are hierarchically clustered with Ward D2 algorithm using Pearson correlation (red-blue color scale) as distance. On the far left, the preservation status of each module is indicated with grey color scale (black-preserved, grey – not preserved). Clusters of modules with concordant annotation are highlighted with dashed squares. On the left, EGs (purple-orange color scale) for the exemplar compounds tunicamycin, cyclosporine A, Acetaminophen, Omeprazole are shown for the 87 modules, in dose- and time-responses. On the far right, modules names are shown together with their main annotation (available for the seed modules), in red highlighted the modules show in table 1.

#### Cluster 1: immune response

Cluster 1 includes all the seed modules annotated for immune response pathway terms (Table 1), except PHH:247 (cluster 8). Add-in module PHH:317, which showed no annotation terms, contains an interleukin receptor (IL18R1), but also some genes induced by inflammatory molecules, for which little is known regarding involvement with immune response (GSAP, CSTO). Similar to the observations obtained with TNF-α and interferon-α, inflammatory modules clusters into two subgroups, one enriched with NF-κB modules, the other in STAT regulated modules.

#### Cluster 5: mitochondria

Cluster 5 contained seed modules annotated with mitochondria related terms, but also add-in modules with less clear annotation. Nevertheless, when looking for protein-protein interactions among genes belonging to this group of modules, high degrees of connections are found for a large subgroup of genes, connecting within modules and showing high corEG (Wilcoxon rank sum test p = 7.72E-07, Figure S4).

#### Cluster 3, 4 and 8: cellular stress responses

The above characterized stress modules, like oxidative stress, ER stress response, heat shock and DDR, clustered in isolated sub-groups within clusters 3, 4 and 8. However, these clusters also included other processes less commonly associated with prototypical stress responses. For example, Cluster 3 contained the seed proteasome module (PHH:76), a module containing two growth arrest and DNA-damage inducible (GADD) genes, *GADD45A* and *GADD45B* (PHH:243), the ATF4-regulated modules (PHH:15 and PHH:295), a module containing several inducible heat shock protein transcripts (PHH:131) and a module containing two NRF2 target genes, *MAFG* and *MXD1* (PHH:337). Cluster 3 also contained two RNA processing modules (PHH:42, PHH:19) associated with the nucleolus/ribosome biogenesis and mRNA processing, respectively. Cluster 8 contains the seed DDR modules (PHH:59 and PHH:83), both of which contain multiple TP53 regulated genes, and a seed mitochondrion module (PHH:2) along with modules enriched for fatty acid/cholesterol metabolism modules (PHH:40, PHH:240) but also cellular homeostasis modules, like PHH:1 (cell cycle), and PHH:11 (metabolism).

### The expanded interaction map describes the interplay between cellular processes in response to chemical treatment

The interplay across module clusters appeared to capture elements of the dynamic interactions among biological response networks (off-diagonal correlation in Figure 3). Clusters can interact positively, e.g. the mitochondria cluster 5 and the RNA processing cluster 6 (which contains both ribosome RNA (rRNA) and mRNA processing modules), or negatively, e.g. activation of stress cluster 3 is commonly associated with down-regulation of modules in cluster 8 (cell cycle and mitochondrion). Interestingly, half the modules, largely those annotating for NF-κB, in immune response cluster 1, showed positive correlation (induced) with activation of processes in clusters 2 and 3 while the other half, including the STAT modules, were inversely correlated and down-regulated. Cluster 1 showed a similar bifurcation with cluster 8 but in the opposing direction, i.e. the NF-κB modules positively correlated with clusters 2 and 3 were negatively correlated with cluster 8 while the STAT modules showed a positive correlation.

An integrated understanding of interaction among complex cellular processes is critical for a mechanism-based risk assessment and requires identifying KE associations with an apical endpoint. Compound with higher versus lower risk may perturb more cellular processes in progressing to cytotoxicity while others may have a more adaptive phenotype. Evidence for progressive perturbation (or lack thereof) can be derived within the module cluster correlation landscape by comparing the module activation by exemplar compounds (Figure 3, right). For example, tunicamycin is a prototypical ER stress inducer and has a low cytotoxic concentration in human hepatocyte cells and a low LD50 of 2 mg/kg in mice (Morin & Bernacki, 1983; S. Zhang et al., 2014). Tunicamycin clearly activates the ER stress response in PHH both via the ATF6 arm in cluster 4a (PHH:13 and 62), the ATF4 arm (cluster 3) and fatty acid metabolism (cluster 8 and cluster 4), concurrently with strong repression of rRNA and mRNA processing (cluster 6) and immune response activation (cluster 1). In contrast, cyclosporine A, which is less toxic with an LD50 in mice ranging from 96-2803 mg/kg (depending on the route of administration (Sax, 1975)), primarily triggered the ATF6-ER response (cluster 4a). Taken together, these observations suggest that tunicamycin is inducing a more severe response impacting diverse cellular biological processes, whereas cells challenged with the used concentrations of cyclosporine A showed limited module perturbations.

A different picture emerged when we considered two prototypical drugs that can induce oxidative stress: acetaminophen and omeprazole. For both, immune responses were primarily repressed (cluster 1), while oxidative stress modules (cluster 4) were activated. In addition, omeprazole activates modules in cluster 3, concurrent with a deactivation of modules in cluster 8. Specific transcription regulation modules are activated, including PHH:4 which contains several stress-induced TFs (ATF4, ATF3, NFE2L2, TP53) and the xenobiotic stress module, including CYP enzymes (cluster 4). Interestingly, PHH:358 (cluster 4) contains CYP1A1 and CYP1A2, canonical AHR targets, and omeprazole is a known AHR ligand (Safe et al., 2020). Taken together, these observations suggest that omeprazole activates oxidative stress, coupled with the activation of processes to constrain protein damage (ATF4, proteasome and heat shock modules in cluster 3) and repression of fatty acid metabolism and cell cycle progress (cluster 8).

These examples illustrate the utility of the PHH TXG-MAPr stress-pathway landscape for mechanistic evaluation of compounds and for comparing compounds with similar modes of action but with different degrees of toxicity.

### Visualizing external datasets in the TXG-MAPr tool reveals good correlation between different cell types and gene expression platforms in the mode of action

Integrating historical and new datasets to derive mechanistic interpretations is important, but it is often hard to determine if merged information, such as WGCNA modules, provides a consistent view of biological responses across experiments. The high variability of gene-level analyses, and the high penalty for multiple comparisons also complicate data integration. To validate the utility of the PHH TXG-MAPr for integrating and comparing external datasets, we created a data upload function, which calculates new module eigengene scores (EGs) from the gene log2FC of the uploaded data and overlays the scores onto the module dendrogram for visualization (for further details, see Material and Methods paragraph Uploading new data files). To further describe CSA mode of action, we processed and uploaded samples from the GEO database of CSA prolonged repeated exposure followed by recovery in collagen sandwich cultured PHH (GSE83958 (Wolters et al., 2016)) into the PHH TXG-MAPr tool (Figure 4A). The uploaded CSA dataset showed strong correlation (Pearson R > 0.8) in module activation and repression among the three timepoints. A heatmap of module EGs (absolute EGs > 2) separates repressed and activated modules in different clusters (Figure 4B). ER stress annotated modules (PHH:13 and PHH:62) and an ATF4 module (PHH:15) are located in the same cluster, which are strongly activated by CSA at high dose levels (Figure 4C). In addition, the 5 day 30 µM CSA treatment with 3 days recovery looks most similar to the CSA_24h_6 µM data from TG-GATEs based on module EGs correlation (R Pearson = 0.67, Figure 4D). The absolute Pearson correlation between module EGs across the uploaded CSA conditions was higher than the absolute correlation between the gene log2FC (Supplementary Figure S5F). These uploaded data confirm that high dose cyclosporine A induces a strong ER stress response amongst others, which is present for several days during daily exposure and persists even after washout. A similar ER stress response (PHH:13 and PHH:62) was also present in HepG2 hepatocytes exposed to CSA (GEO dataset: GSE45635 (Van den Hof et al., 2015a)) and ER stress module activation was seen at low (3 µM) and high (20 µM) dose (Suppl Figure S5A, S6 cluster 5 red line).

**Figure 4.**
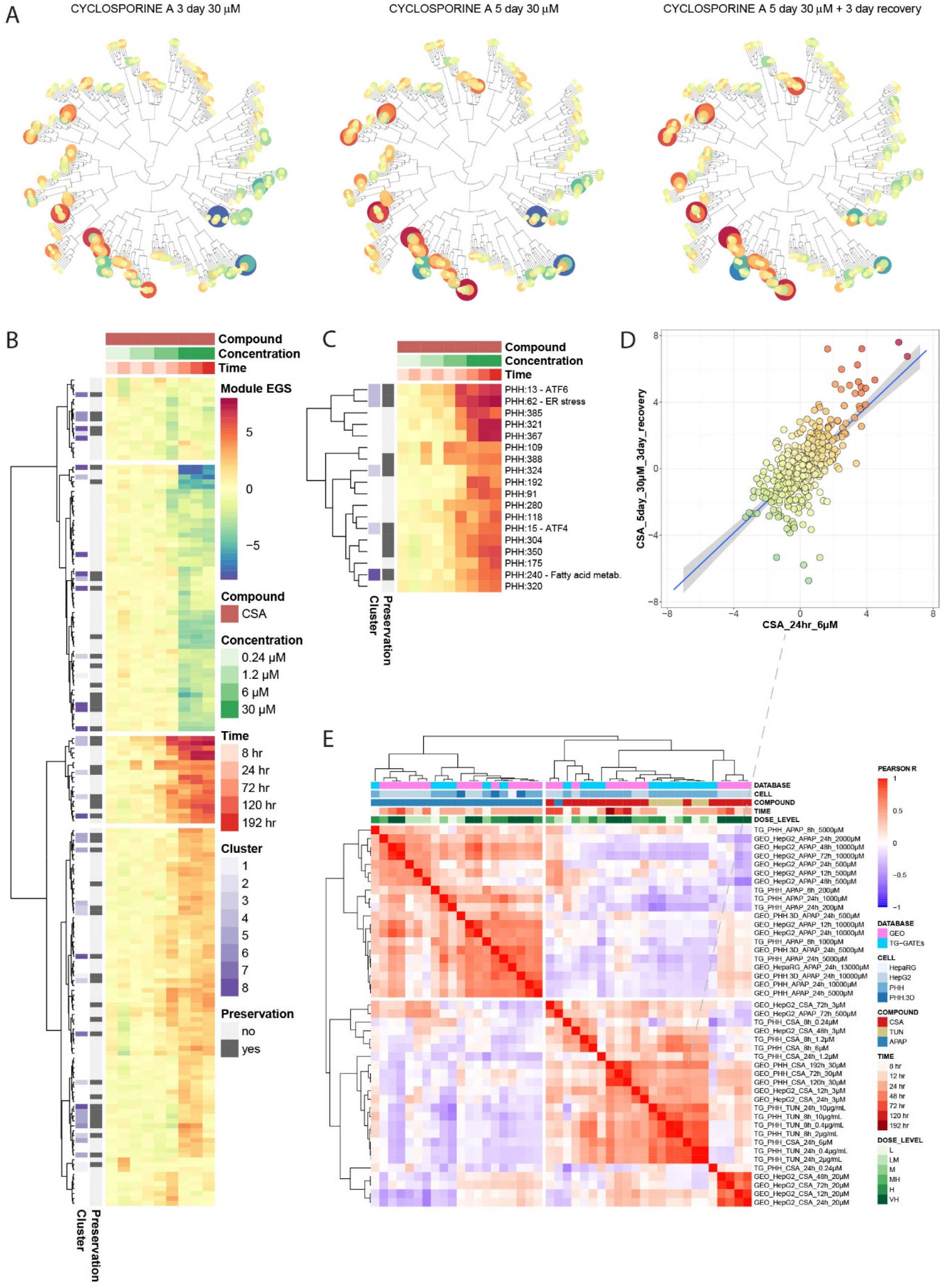
Upload external data into the PHH TXG-MAPr. A) Phylogenic tree view of module EGs upon 30 µM cyclosporine A exposure at 3 day (left), 5 day (middle) and 5 day + 3 day recovery (right). B) Heatmap of all modules that have at least one condition with absolute EG score > 2 for cyclosporine A treatment. All concentrations and time points from TG-GATEs (0.24, 1.2 and 6 µM) and uploaded GEO: GSE83958 dataset (30 µM) are shown. Modules are clustered by Euclidean distance, Ward.D2 method. The purple color scale indicates to which cluster of Figure3 each module belongs, and grey color scale indicates the preservation status. C) Zoom in for the third cluster obtained from the heatmap in B. Modules that have a strong annotation for ER stress (PHH:13 and PHH:62) or ISR (PHH:15) are in this module cluster and are strongly induced by 6 - 30 µM cyclosporine A. C) Compound correlation plot comparing the module EG scores of the uploaded 30 µM CSA data at 5 day + 3 day recovery (y-axis), with CSA_24hr_6µM (x-axis) available in the PHH TXG-MAPr (Pearson R = 0.67). E) Cluster correlation heatmap showing the Pearson R correlation between the conditions in TG-GATEs data (CSA, TUN, APAP) and the uploaded datasets with CSA and APAP exposures in PHH, HepG2 and HepaRG cells at various time points (see labels). Compounds cluster by mode-of-action, because ER stress inducers TUN and CSA are clustering together and show low correlation with the oxidative stress inducer APAP.

Additional datasets of human hepatocytes (PHH, HepG2 and HepaRG) exposed the oxidative stress inducer acetaminophen (APAP) were analyzed to investigate the application of the PHH TXG-MAPr for a different compound (Suppl. Figure S5B-E). The APAP datasets show a clear oxidative stress response (PHH:144, PHH:325) as well as many other module perturbations, which is independent of the cell type or microarray platform (Figure S6, cluster 8 red line). Pearson correlations between CSA and APAP treatments in TG-GATEs and the uploaded data from GEO are shown in a cluster correlation plot (Figure 4E). Hierarchical clustering based on correlation distance between samples separates the data based on the mode of action, clustering ER stress inducers cyclosporine A and tunicamycin together and separated from oxidative stress inducer acetaminophen. In addition, different hepatocyte cell cultures (3D PHH, HepG2 and HepaRG) show good correlation with the same compound exposure, suggesting that a similar mode of action can be captured by other hepatocyte cells. GEO data of PHH exposed to APAP that was analyzed on two different Agilent microarray platforms correlate well with the TG-GATEs data at the same concentration (GEO_PHH_APAP_24h_5mM / GEO_PHH.3D_APAP_24h_5mM versus TG_PHH_APAP_24h_5mM), even though the average module coverage was 84% or 93% for the Agilent chips (e.g. only 84% or 93% of genes per module were measured in the chip). This indicates that the TXG-MAPr supports the analyses of gene expression data from different platforms/chipsets. In contrast, low dose or early time-points do not correlate well with the same compound, likely due to the minimal module perturbations in these conditions (data not shown).

Overall, the PHH TXG-MAPr upload and analysis functions are powerful tools to compare both global biological responses across disparate datasets and TG-GATEs data, while identifying specific modes-of-action, even for other hepatocyte cell types or other microarray platforms, suggesting that co-expression analysis can obviate technical issues when comparing across expression datasets or platforms.

### Leveraging WGCNA to interpret targeted RNA Seq datasets

Although high throughput technologies allow many samples to be processed in parallel, the high costs of whole transcriptome profiling has led to the use of targeted RNA sequencing approaches which are gaining popularity for toxicology studies (Mav et al., 2018). We reasoned that mapping a targeted gene sets to co-expression modules might yield sufficient data to impute more detailed and biologically relevant information. We tested this concept using the PHH TXG-MAPr by uploading a targeted RNA Seq data set, TempO-Seq, obtained from treating plated cryopreserved PHH from 50 different human donors in a detailed dose-response protocol with tunicamycin at two time points (0.0001 to 10 µM, 8 and 24 hours, Niemeijer et al. 2021). We calculated the modules EGs for each donor-concentration-time point combination, a total of 600 different conditions, following the approach detailed in Material and Methods section.

Of the 2708 genes measured in the TempO-Seq S1500+ gene set, only 1830 genes are mapped to the PHH modules (Supplementary Table S9). Genes not mapping were excluded from the PHH network during modules generation because they did not have robust co-expression patterns and are included in the PHH TG-GATEs ‘gray’ module (Langfelder & Horvath, 2008). Although some modules were not covered by any genes from the S1500+ set (coverage of 0%), modules selected in Figure 3 have significantly higher coverage (Wilcoxon test, greater alternative, p<0.05 and Figure 5A and B), suggesting that the preserved and well annotated modules are better represented in the S1500+ set. In addition, the coverage was higher when the hub gene was present in the S1500+ set (Figure 5B). EG calculation depends on both the fold change and the correlation with the eigengene score of each gene (corEG, a measure of hubness), therefore we assessed the corEG values for S1500+ genes as a gene quality metric. At the module level, the average corEG does not seem to be heavily impacted by the subsampling of gene space, and modules in the expanded seed set (Figure 3) show higher concordance (Figure 5C, Pearson R 0.84 versus Pearson R 0.65 including all modules). Additionally, the S1500+ set shows higher average corEG compared with random draws of genes from PHH module (adj. p-value < 0.05, see Material and Methods Data analysis section). We also assessed coverage within the set of the preserved modules; only two show coverage of 0%, PHH:103 and PHH:105 (Supplementary Table S10). Thus, the S1500+ set represented a set of genes that overlap, to a greater or lesser extent, nearly all of the preserved modules and higher quality relative to their corEG values.

**Figure 5.**
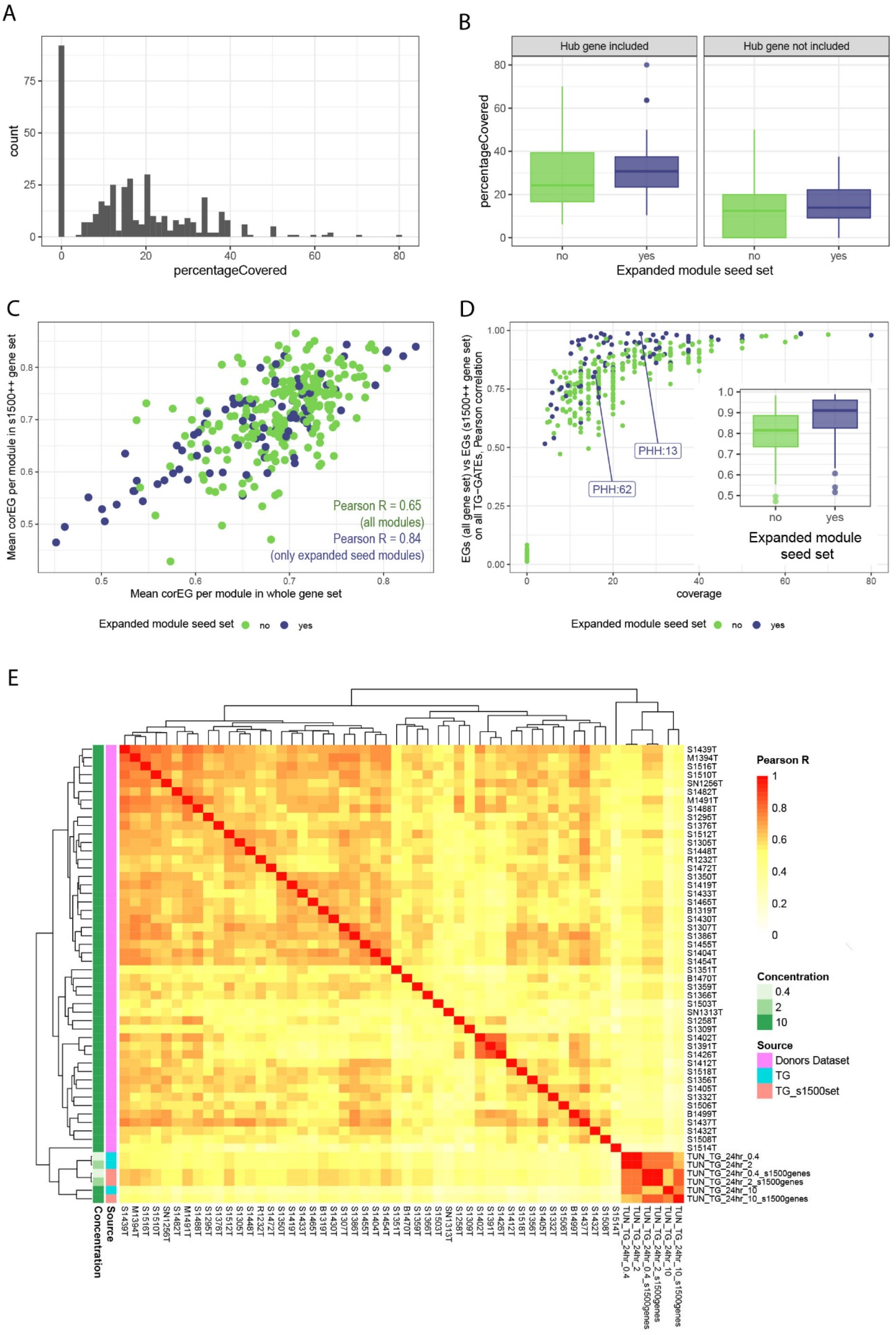
Upload of 50 donors PHH study S1500+ TempO-Seq set. A) Histogram of modules coverage when uploading the targeted TempO-Seq gene set. Frequency (y axis) of modules percentage covered by the uploaded targeted gene set (x axis). B) Percentage covered (y axis) for PHH modules, grouped on the x axis by whether they are part of the correlation matrix in Figure 3 (blue) or not (green). The plot is divided into two sections, the first one showing modules where the hub gene was included in the uploaded gene set, the second one showing modules where the hub gene was not included in the uploaded gene set. C) Mean correlation Eigengene for each module, whole genes set (x axis) versus after upload with the targeted TempO-Seq gene set (y axis). Points are colored based on whether they are part of the correlation matrix in Figure 3 (blue) or not (green). D) Pearson R correlation for each module between EGs calculated with the complete gene set, and EGs calculated with the S1500+ gene set, for all TG-GATEs experiments. Points are plotted against module coverage (x-axis) and are colored based on whether they are part of the correlation matrix in Figure 3 (blue) or not (green). Inside: Pearson correlation calculated as previously grouped by whether they are part of the correlation matrix in Figure 3 (blue) or not (green). Modules with coverage = 0% had been excluded. E) Cluster correlation heatmap (complete clustering, Euclidean distance) showing the Pearson R correlation between modules EGs of the 50 donors dataset samples (10 µM, 24 hr, pink color code on the left), TG-GATEs tunicamycin data obtained with the complete gene set (aquamarine color code on the left), and TG-GATEs tunicamycin data using only the S1500+ gene space (salmon color code on the left).

To test the impact of calculating EGs based on the S1500+ gene set, we recalculated EGs for all TG-GATEs samples using only the reduced S1500+ gene set and calculated the Pearson correlation per module compared to EGs calculated using the complete gene set (Figure 5D). Modules even with minimal, but above zero, gene coverage show good EGs correlation (always higher than 0.5), and modules in the expanded seed set (Figure 3) show on average higher correlation (Wilcoxon test, p value of 2.72e-09). Using the ER stressor tunicamycin as a model compound, we performed a cluster correlation analysis using EGs for TG-GATE PHH data, calculated using both whole chip or just the S1500+ genes, and compared results with a set of TempO-Seq, S1500+ data derived from 50 donor PHH cultures treated with tunicamycin (Figure 5E, Supplementary Figure 8). The Affymetrix- and S1500+ derived scores for TG-GATEs tunicamycin treatments were always >0.7 and clustered together. However, there was considerable variation within the donor set at the highest dose (10 µM). A subset of donors shows higher similarity within themselves and with TG-GATEs tunicamycin samples, especially for medium concentration and when calculated with the S1500+ gene set (Figure 5E, donors in upper left quadrant, to compared to TG-GATEs sample in the right bottom quadrant). Another subset of donors showed lower similarities within each other and lower similarity with TG-GATEs tunicamycin samples (Figure 5E, lower right quadrant).

Thus, a targeted gene set can be used as input to derive EGs for modules derived from the whole transcriptome and impute biological responses on EGs.

### Assessing donor variability captured by PHH modules

Since modules with similar biological annotations cluster together based on EGs scores (Figure 3) we reasoned that identifying sources of variability at the level of clusters might be more robust than at the individual module level. Therefore, we clustered modules based on the S1500+ EGs of the 50-donor dataset similarly to Figure 3, and assessed the overlap between clusters (Figure S9). Some, but not all, modules clusters derived from the donor data show significant overlap with the groups obtained using the entire TG-GATEs set in Figure 3 (Figure S9B, exact Fisher p-values and numerical overlaps are shown). Particularly, RNA processing, stress clusters (ATF4 and ATF6) and transcriptions regulation modules are clustering similarly. The modest overlap is not surprising since the donor dataset is derived with a single compound with the PHH donors as the primary source of variability (Figure 5E).

We then made use of modules EGs of S1500+ clusters 1-8 to study donor variability in response to tunicamycin. Donor variability can be attributable to genomic makeup, but also lifestyle, disease and age status of each individual. Consequently, we analyzed relationships between module scores, reflecting induction or repression of underlying genes, and the presence (positives) or absence (negatives) of donors’ traits, i.e. sex, the presence of cancer, liver pathology, hypertension, diabetes, smoking habit (Supplementary Table S11).

We applied the methodology described in (J. J. Sutherland et al., 2018) and averaged modules EGs within each module clusters and modelling each concentration separately. We calculated Cohen’s d effect sizes (*eff*, (Cohen, 2013)) and performed logistic regression treating avgAbsEG (average absolute cluster score) as a covariate and quantified the p-value for each module in explaining the odds of toxicity (*signed_log10_p_adj*) reflecting the adjustment for differences in overall gene expression, i.e. avgAbsEG (Supplementary Table S12). Next, we calculated the same relationship for individual modules EGs (Supplementary Table S13).

Following this approach, we were able to identify modules and the underlying biological processes, which are uniquely associated with a donor’s trait upon tunicamycin exposure, and with increasing association levels in a dose-response relationship (Figure 6). Module clusters have variable number of genes included but still show good gene quality as quantified by the high average correlation with EGs calculated with the entire TG-GATEs gene set, especially compared to modules excluded in these clusters (Figure 6 and Figure 5D). We focused on the presence of liver pathology, that influences donor’s response based on different processes both positively and negatively (Figure 6A). Modules annotated for DDR (PHH:83), protein folding (PHH:95) and oxidative stress (PHH:325 and PHH:144) are positively associated (induced) with liver pathology phenotype of the donor (Figure 6A, Supplementary Table S12, Supplementary Figure S10). Modules annotated for immune response had a negative association (repressed) with liver pathologies (PHH:12, PHH:26, PHH:44, PHH:22, Figure 6A, Supplementary Table S12 and S8 for module annotation). In particular, donors with liver pathologies show lower EGs for these modules, corresponding to lower log2FC values of the genes belonging to the modules, all annotated for being involved in inflammation and immune response (Figure 6B and Supplementary Figure S10C, showing genes in module PHH:12 with corEG > 0.8). To a closer inspection, genes with lower log2FCs are the result of already higher basal expression levels in PHH donors with pre-existing liver pathology (Figure 6D, showing normalized counts for PHH:12 genes with significant difference between donors with or without liver pathologies, only with DMSO treatment, Wilcoxon test, adj. p-value < 0.05).

**Figure 6.**
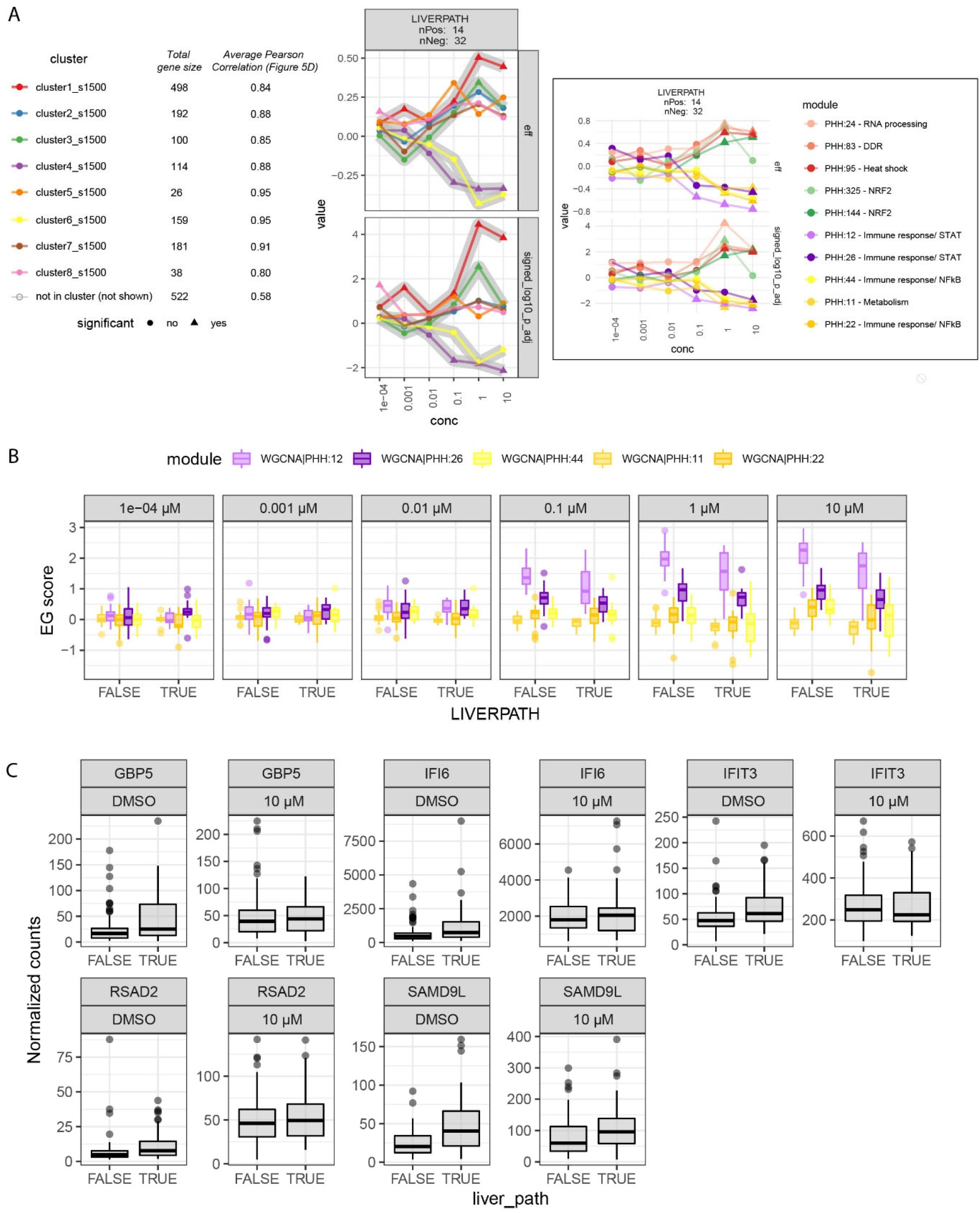
Some PHH modules are associated with donor’s traits. A) Dose-response plot of effect size and adjusted p value (log10 transformation, keeping the sign) resulting from the association of modules clusters to donors’ presence of liver pathologies. Modules clusters indicated by different color, number of donors showing (positive) and not showing (negative) the indicated trait are shown in the plot label. On the right, effect size and adjusted p value (log10 transformation, keeping the sign) of individual modules to trait associations most prominently contributing to the overall cluster associations highlighted with grey shadows. Modules are colored in different saturation level of the same hue of the cluster they belong to. B) Boxplot of modules’ EGs negatively associated with liverpath, grouped by increasing concentrations and presence/absence of liver pathologies. C) Boxplot of normalized counts of genes belonging to module PHH:12 and significantly different between donors with or without liver pathologies with only DMSO treatment (Wilcoxon test, BH adj. p-value < 0.05).

To conclude, pre-existing traits of PHH donors influence the cellular response to an ER stressor (tunicamycin). Such an influence can be attributable not to different magnitude of ER stress response activation, but to variations in the activation of accompanying processes like immune response or oxidative stress.

## Discussion

In this work, we presented a novel Shiny web tool that allows users to apply a gene co-expression network (WGCNA) approach to investigate toxicity mechanisms rapidly and in the context of biological responses preserved between human and animals. This unbiased approach is not weighted toward current knowledge and can reveal unknown relationships between genes and phenotype data, but it also encompasses known biological response networks through modules annotation. In addition, this approach is tailored for a gold standard in vitro testing system: primary human hepatocytes (PHH).

Toxicogenomics data can be difficult to interpret due to its high dimensionality and possible low signal-to-noise ratio (Ideker et al., 2011). We reduced gene expression dimensionality from 10^4^ to 10^2^ by taking a modular approach and constructing networks of co-expressed genes from a large PHH transcriptomic dataset (Igarashi et al., 2015). A set of 398 modules are presented in an intuitive visualization format, the PHH TXG-MAPr. A simplified upload function enables calculation of eigengene scores (EGs) from external sources, allowing new data to be integrated and interpreted in the PHH TXG-MAPr analysis environment which contains 158 chemical comparators combined in 941 different treatment conditions. Pathway enrichment and TF associations contribute to mechanistic interpretation of modules, allowing the user to identify key processes that are different and in common between compounds responses. This represents a starting point for mechanistic interpretation across risk assessment applications, for example, to derive data-driven quantitative AOPs based on MIE and KE represented by module EGs. Stress induced pathways interact in cascades of perturbations, balancing adaptive and progressive responses. We showed how modules and correlation between them can represent connected mechanisms (Figure 3) and used these to describe compounds’ mode of action. Although correlation analysis is not sufficient to establish causality, it can suggest potential areas for additional analysis in order to define cause-and-effect relationship, e.g. through validation of a mechanism-based risk assessment framework (Liu et al., 2019). Additionally, gene co-expression modules allows to understand how applicable a pathway is to other species (Perkins et al., 2015).

One of the most vexing problems in risk assessment is indeed understanding whether toxicology data are likely to extrapolate across species, e.g., from rodent to human, or from one model to another, e.g. from *in vitro* to *in vivo*. Co-expression analysis formalizes methods that allow comparison of node-edge relationships at the gene level (Langfelder et al., 2011). As a result, we were able to define a subset of modules, and the associated biological themes, with higher preservation between *in vitro* human and rat hepatocytes models. Interestingly, a number of preserved PHH modules show also overlap with published signatures of rat xenobiotic receptor activation (Podtelezhnikov et al., 2020): to mention a few, PHH:358 perfectly overlaps with the AHR signature, PHH:16 shows very high enrichment with gene members of the SREBP signature and prominent annotation for cholesterol biogenesis (Supplementary Table S9). A remarkable observation of our results was the notable difference in preservation for some toxicologically relevant and well characterized stress pathways. For example, ER stress response is highly preserved, while DNA damage (p53-driven gene expression) and oxidative stress (Nrf2-driven) responses are not. Interestingly, when comparing module preservation results, we found that *in vivo* rat liver and RPH show higher similarities (Pearson R 0.85 and 0.60 for Z-summary and Median Rank, respectively). In addition, Nrf2/p53 modules show good preservation between PHH and HepG2 data (data not shown), suggesting the differences in module preservation are influenced by species rather than cultured conditions. We expect that preservation results depend, to some extent, on the input dataset composition. For instance, how well a certain process is captured in gene expression data from the input experiments, which can influence gene-to-gene correlation patterns, could be a hypothesis to be explored in future studies. Nevertheless, literature reports that kinetics and activation of the non-preserved Nrf2/p53 processes are different across species (MacRae et al., 2015; Martin & Chang, 2018; Monroe et al., 2020), suggesting that preservation of co-expression networks is a promising approach for the evaluation of the applicability domain of new approach methodologies (NAMs) for extrapolating safety assessments across species (Parish et al., 2020).

Comparison with external datasets is a crucial application of the PHH TXG-MAPr tool. Gene level comparisons can be risky given the likelihood of variability across models and the low signal-to-noise ratio. Within TG-GATEs data, we showed that compound correlations are improved at the module level (Figure 1C, S1C, S5F). Additionally, module EGs derived from microarray-based external GEO data shows high similarity across external and TG-GATEs datasets for the same compounds, suggesting that gene-level noise is dampened when results are evaluated at the modules level. Using the external GEO datasets, we also confirmed that ER stress is an early event in the mode of action of CSA in different liver cell types. In contrast, CSA did not show a good correlation with the MoA of APAP datasets, suggesting that oxidative stress is a secondary/late effect to high dose CSA exposure. Although several publications indicate that oxidative stress is involved during CSA induced cholestasis in liver (Kawamoto et al., 2017; Koido et al., 2020; Wolters et al., 2016), our analysis of CSA exposed hepatocytes using the TXG-MAPr tool showed that the ER stress should not be neglected in the mechanism of CSA induced liver injury as was also suggested by other literature (Koido et al., 2020; Van den Hof et al., 2015b). To validate the applicability of the PHH TXG-MAPr for other transcriptomic platforms, we showed that Agilent microarray data of APAP exposure in PHH showed strong correlation with Affymetrix based data in TG-GATEs.

Integrating and comparing data across platforms represent a great challenge, especially in cases where targeted gene sets have been designed to be representative for the whole transcriptome (Mav et al., 2018; Soufan et al., 2019). A reduced number of features is convenient to diminish complexity and costs but can weaken the power of enrichment analysis given the smaller gene set background. Imputing the entire transcriptome response prior to analysis is one solution to this problem (Mav et al., 2020), but may not be tailored for the testing system in use. Using the TempO-Seq S1500+ gene set platform, we showed that EGs calculated with PHH co-expression modules can impute biological response patterns directly when analyzing and interpreting PHH data from targeted gene sets in the context of whole transcriptomic data. Although the targeted gene space is dramatically reduced compared to whole transcriptome data in TG-GATEs (∼17%, 1830/10275 genes), the modules included in our collection of mechanistic relevant modules (Figure 3) had significantly higher coverage compared to the entire module set, and include genes of high quality (as quantified by the corEG). EGs scores obtained with the complete gene set and with the S1500+ set are in good agreement, and the mechanistic relevant modules had significantly higher agreement than the rest. Taken together, these findings suggest that results based on only genes in the S1500+ platform can be extrapolated to recapitulate the whole transcriptome modules score variability.

Following this strategy, we circumvented the limited gene coverage of the S1500+ set by leveraging the correlation structure of modules EGs to define biological responses relevant to the differential sensitivity across responses of 50 donor hepatocyte treated with an ER stress inducer, tunicamycin. Donors’ traits, particularly the presence of liver pathologies, show significant dose-response related associations with clusters of modules referring to specific biological responses, namely immune, DNA damage, oxidative stress responses and protein folding. The presence of liver pathologies seems to be associated with a weakened activation of innate immune response upon ER stress induction. Interestingly, genes involved in innate immunity in donors with pre-existing liver pathologies show already higher expression values without treatment compared to donors without liver pathologies. Under normal conditions, if there is excessive stress, ER stress evokes inflammatory responses via the activation of the three UPR branches, which may sensitize to adverse events (Duvigneau et al., 2019; Fredriksson et al., 2014; Peng et al., 2020). Donors with pre-existing liver diseases show already activated inflammation, therefore possibly resulting in even higher sensitivity. Modules related to DNA damage response, protein folding and oxidative stress response appear to be mildly sensitized in patients with liver pathologies upon ER stress induction. That is in accordance with a condition of chronic stress and continuous attempt to restore homeostasis (García-Ruiz & Fernández-Checa, 2018). Our results partly overlap with findings in recent GWAS DILI studies, where cholestatic injury phenotype has been associated with UPR and mitochondrial stress, but most importantly with an increased sensitivity to ROS upon additional drug treatment (Koido et al., 2020). Interestingly, ER stress response did not show a connection with donor traits: ER stress related modules, collectively, were a stable and preserved set of biomarkers for the unfolded protein response. Ultimately these findings highlight how pre-existing liver diseases can be a confounding factor when interpreting data from *in vitro* models using donor hepatocytes and that conserve alterations even in 2D culture conditions.

In conclusion, the TXG-MAPr takes advantage of the modular nature of gene co-expression networks to achieve mechanistic relevant, cross-species and cross-platform evaluation of toxicogenomic data. By using the PHH TXG-MAPr, we were able to reduce toxicogenomic data dimensionality to an interpretable set of mechanistically relevant groups of co-expressed genes. The network-based structure of the TXG-MAPr allows one to evaluate the similarity of such components in other experimental systems and species. Therefore, preserved modules between primary human cells and rat systems could be defined, which could help with interspecies translation during risk assessment. Finally, we illustrated how data from multiple sources, including data from targeted gene sets, are applicable to upload in the TXG-MAPr, as well as how the approach can be applied to shed new light into donor-to-donor variability in a high throughput transcriptomic setting. Overall, we demonstrated that gene co-expression analysis coupled to a facile visualization environment, the PHH TXG-MAPr, is a promising approach to analyze in vitro human transcriptomic data and derive mechanistic interpretation, and therefore a substantial step forward towards integration of transcriptomic data in mechanistic risk assessment practices.

## Abbreviations

APAP: acetaminophen
avgAbsEG: average absolute eigengene score
corEG: correlation eigengene score
CSA: cyclosporine A
DILI: drug?induced liver injury
DDR: DNA damage response
EGs: eigengene score
ER: Endoplasmic Reticulum
GO: gene ontology
ISR: integrated stress response
MoA: Mode of Action
PHH: primary human hepatocytes
RPH: rat primary hepatocytes
TF: transcription factor
TG: TG?GATEs
TUN: tunicamycin
TXG: toxicogenomic
WGCNA: weighted gene co?expression network analysis

## Acknowledgements

This work was supported by the EU-EFPIA Innovative Medicines Initiative 2 (IMI2) Joint Undertaking TransQST project (grant number 116030) and eTRANSAFE project (grant number 777365) (this Joint Undertaking receives support from the European Union’s Horizon 2020 research and innovation program and EFPIA), the EC Horizon2020 EU-ToxRisk project (grant number 681002), and the Cosmetics Europe/CEFIC Liver Ontology project.

## Supplementary Figure captions

**Figure S1. Module dendrograms construction**. A) Module dendrogram is shown together with the anti-clockwise labelling system of branches. B) Module dendrogram is shown highlighting manual adjustment of branch lengths C) Boxplot of Pearson correlation between different conditions. TG-GATEs conditions have been grouped by compound, and Pearson correlation calculated between different treatment conditions in each group (the same compound at different concentrations/time points). The maximum Pearson correlation per compound is plotted when considering correlation calculated with all log2FC genes, log2FC of the genes included in the PHH modules, or modules EGs.

**Figure S2. Module dendrograms over time and concentrations and for different compounds in TG-GATEs at high concentration, 24hrs**. A) cyclosporine A module dendrograms at increasing time points (from left to right) and increasing concentrations (from top to bottom). 2hr time point has not been measured for this compound. B) Acetaminophen module dendrograms at increasing time points (from left to right) and increasing concentrations (from top to bottom). 2hr time point has been measured for this compound. C) Module dendrograms for 5 compounds annotated as “MostDILIConcern” according to the DILIRank classification. D) Module dendrograms for 5 compounds annotated as “LessDILIConcern” according to the DILIRank classification. E) Module dendrograms for 2 compounds annotated as “noDILIConcern” according to the DILIRank classification. F) Module dendrograms for 3 compounds not annotated in the DILIRank classification.

**Figure S3. Z summary and Median Rank in the module dendrogram view**. A and B) Z summary preservation scores of PHH modules in Rat in vivo Liver data TG-GATEs (A) and in Rat Primary Hepatocytes (RPH) TG-GATEs (B), shown in the module dendrogram view. All modules are colored according to their Z summary statistic (range from blue to red, being blue Z-summary ∼0 and deep red Z-summary ∼30). Size of dots representing the modules is proportional to Z-summary. C and D) Median Rank preservation scores of PHH modules in Rat in vivo Liver data TG-GATEs (C) and in Rat Primary Hepatocytes (RPH) TG-GATEs (D), shown in the module dendrogram view. All modules are colored according to their Median Rank statistic (range from blue to red, being blue MedianRank = 396 and deep red MedianRank = 1). The modules having Median Rank <100 are shown with dots having double the size.

**Figure S4. Protein-Protein Interaction plots of genes in the mitochondrial cluster**. A) Protein-Protein Interaction plot of genes belonging to modules part of the mitochondrial cluster (Figure 3) have been generated with String-DB and modified in Cytoscape. Nodes (genes) are colored for module membership. Edges lengths are proportional to the combined interaction score (http://version10.string-db.org/help/faq/). B) Same plot, each node is colored for the corresponding gene correlation EG score. C) Boxplot of absolute gene corEG for the nodes connected in the bigger cluster in plots A and B, or not connected.

**Figure S5. TXG-MAP dendrograms of the uploaded conditions into the PHH TXG-MAPr**. A) GEO GSE45635: HepG2 exposed to 3 and 20 µM CSA at 4 time points (12, 24, 48 and 72 h). B) GEO GSE53216: HepG2 exposed to 0.5 and 10 mM APAP at 4 time points (12, 24, 48 and 72 h). C) GEO GSE74000: HepG2 and HepaRG, exposed to IC10 concentrations of APAP for 24 hours. HepG2 = 2mM; HepaRG = 13mM. D) GEO GSE13430: PHH exposed to 5 and 10 mM APAP for 24 hours. Microarray analysis was done on the Agilent Human 1A Microarray (V2) G4110B platform and shows 84% module coverage when uploading to the PHH TXG-MAPr. E) GEO GSE104601: PHH in 3D co-culture with Kupffer cells exposed to 0.5, 5 and 10 mM APAP for 24 hour. Microarray analysis was done on the Agilent SurePrint G3 Human GE v2 8×60K Microarray platform and shows 93% module coverage when uploading to the PHH TXG-MAPr. All other GEO datasets are analyzed with the Affymetrix Human U133 plus 2.0 array and show 100% module coverage. F) Absolute Pearson correlation between the uploaded PHH datasets for CSA (GSE83958) and the 24 hr CSA HI data in TG-GATEs, using log2FC of all genes, log2FC of genes in modules, or module EGs.

**Figure S6. Heatmap of the uploaded conditions into the PHH TXG-MAPr**. Left, heatmap of all modules that have at least one condition with EG score > 2 for conditions in TG-GATEs data (CSA and APAP) and the uploaded datasets with CSA and APAP exposures in PHH, HepG2 and HepaRG cells at various dose levels and time points (see labels). Modules are clustered by Euclidean distance, Ward.D2 method using pheatmap package in R. Module annotation is shown in the rows of the heatmap. Right, zoom in for cluster 5-8 obtained from the left heatmap. ER stress and ATF annotated modules (PHH:13, 15 and 62) can be found in cluster 5 and are induced by cyclosporine A in PHH and HepG2 at higher dose levels. NRF2 annotated modules (PHH:144 and 325) can be found in cluster 8 and are induced by acetaminophen in PHH and HepG2 at higher dose levels.

**Figure S7. Upload of 50 donors PHH study S1500+ TempO-Seq set**. A) Z-summary preservation score to Rat in vivo Liver (TG) of PHH modules (y axis) plotted against the percentage covered after uploading of the targeted TempO-Seq gene set (x axis). Modules having z-summary score higher than 2, no uploaded genes (percentageCovered = 0) and part of the correlation matrix in Figure 3 (blue) are labelled. B) Median Rank preservation score to Rat in vivo Liver (TG) of PHH modules (y axis) plotted against the percentage covered after uploading of the targeted TempO-Seq gene set (x axis). Modules having Median Rank score lower than 100, no uploaded genes (percentageCovered = 0) and part of the correlation matrix in Figure 3 (blue) are labelled. C) Z-summary preservation score to RPH (TG) of PHH modules (y axis) plotted against the percentage covered after uploading of the targeted TempO-Seq gene set (x axis). Modules having z-summary score higher than 2, no uploaded genes (percentageCovered = 0) and part of the correlation matrix in Figure 3 (blue) are labelled. D) Median Rank preservation score to RPH (TG) of PHH modules (y axis) plotted against the percentage covered after uploading of the targeted TempO-Seq gene set (x axis). Modules having Median Rank score lower than 100, no uploaded genes (percentageCovered = 0) and part of the correlation matrix in Figure 3 (blue) are labelled. E) Correlation EG distribution for all genes in the PHH modules. F) Coverage vs module size. G and H) Distribution of the correlation Eigengene for each module, S1500+ set (G) and original gene set (H, PHH WGCNA modules gene space). Modules are ordered on the x axis by module size (from larger to smaller module) and on the y axis boxplots for each individual module show the distribution of the correlation eigengene. Boxplots are colored based on whether they are part of the correlation matrix in Figure 3 (blue) or not (green).

**Figure S8. Distribution of modules scores for the 50 donors uploaded dataset (all modules)**. A) Boxplot of EGs of modules for the 50 donors uploaded dataset. The first sample (TUN) correspond to TG-GATEs EGs of tunicamycin treated sample B) PCA plots colored by time (left) and concentration (right), using the entire dataset (first row), only high concentration or 24 hr samples (second row) or only low concentration or 8hr samples (last row).

**Figure S9. 50 donors’ modules correlations**. A) Cluster correlation matrix of the 87 modules in Figure 3. Modules are hierarchically clustered with Ward D2 algorithm using Pearson correlation (red-blue color scale) as distance. On the left, the preservation status of each module is indicated with grey color scale (black-preserved, grey – not preserved). Clusters of modules are labelled as they appear in Figure 6. B) Matrix of overlap between module clusters obtained with TG-GATEs dataset (rows) and with the 50-donors dataset (column). In each cell, Fisher exact p value of overlap significance and numerical overlap are shown. Cells are colored proportionally to the p value (white, p value equal to 1, dark green p values < 0.05).

**Figure S10. Module to donors’ traits associations**. A) Dose-response plot of effect size and adjusted p value (log10 transformation, keeping the sign) resulting from the association of modules clusters to donors’ traits. Plots are divided by different traits and modules clusters indicated by different color. Number of donors showing (positive) and not showing (negative) the indicated trait are shown in the plot label. B) Boxplot of log2FC values of genes belonging to module PHH:12 and showing corEG > 0.8. C) Boxplot of log2FC values of genes belonging to negatively associated modules to liver path (besides PHH:12) and showing corEG > 0.7. D) Boxplot of EGs of modules positively associated with liver pathologies, grouped by increasing concentrations and presence/absence of liver pathologies. E) Boxplot of log2FC values of genes belonging to the positively associated modules to liver path and showing corEG > 0.7.

## Notes

https://txg-mapr.eu/WGCNA_PHH/TGGATEs_PHH/

### Competing Interest Statement

The authors have declared no competing interest.

## References

Alexa, A., & Rahnenführer, J. (2007). Gene set enrichment analysis with topGO. R package, 27.

Alvarez, M. J., Shen, Y., Giorgi, F. M., Lachmann, A., Ding, B. B., Hilda Ye, B., & Califano, A. (2016). Functional characterization of somatic mutations in cancer using network-based inference of protein activity. Nature Genetics, 48(8), 838–847. https://doi.org/10.1038/ng.3593

Bailey, J., & Balls, M. (2019). Recent efforts to elucidate the scientific validity of animal-based drug tests by the pharmaceutical industry, pro-testing lobby groups, and animal welfare organisations. BMC Medical Ethics, 20(1), 1–7. https://doi.org/10.1186/s12910-019-0352-3

Barel, G., & Herwig, R. (2018). Network and pathway analysis of toxicogenomics data. Frontiers in Genetics, 9(OCT). https://doi.org/10.3389/fgene.2018.00484

Björnsson, E. S. (2019). Global Epidemiology of Drug-Induced Liver Injury (DILI). Current Hepatology Reports, 18(3), 274–279. https://doi.org/10.1007/s11901-019-00475-z

Breiden, B., & Sandhoff, K. (2019). Emerging mechanisms of drug-induced phospholipidosis. In Biological Chemistry (Vol. 401, Issue 1, pp. 31–46). De Gruyter. https://doi.org/10.1515/hsz-2019-0270

Campos, G., Schmidt-Heck, W., De Smedt, J., Widera, A., Ghallab, A., Pütter, L., González, D., Edlund, K., Cadenas, C., Marchan, R., Guthke, R., Verfaillie, C., Hetz, C., Sachinidis, A., Braeuning, A., Schwarz, M., Weiß, T. S., Banhart, B. K., Hoek, J., … Godoy, P. (2020). Inflammation-associated suppression of metabolic gene networks in acute and chronic liver disease. Archives of Toxicology, 94(1), 205–217. https://doi.org/10.1007/s00204-019-02630-3

Clark, M., & Steger-Hartmann, T. (2018). A big data approach to the concordance of the toxicity of pharmaceuticals in animals and humans. Regulatory Toxicology and Pharmacology, 96, 94–105. https://doi.org/10.1016/j.yrtph.2018.04.018

Cohen, J. (2013). Statistical Power Analysis for the Behavioral Sciences. In N. Lawrence Erlbaum Associates: Hillsdale (Ed.), Statistical Power Analysis for the Behavioral Sciences. https://doi.org/10.4324/9780203771587

Colombo, M., La Vecchia, C., Lotti, M., Lucena, M. I., Stove, C., & Paradis, V. (2019). EASL Clinical Practice Guideline: Occupational liver diseases. In Journal of Hepatology (Vol. 71, Issue 5). https://doi.org/10.1016/j.jhep.2019.08.008

Csardi, G., & Nepusz, T. (2006). The igraph software package for complex network research. InterJournal Complex Systems, Complex Sy(1695), 1695. http://igraph.sf.net

Duvigneau, J. C., Luís, A., Gorman, A. M., Samali, A., Kaltenecker, D., Moriggl, R., & Kozlov, A. V. (2019). Crosstalk between inflammatory mediators and endoplasmic reticulum stress in liver diseases. Cytokine, 124, 154577. https://doi.org/10.1016/j.cyto.2018.10.018

Foufelle, F., & Fromenty, B. (2016). Role of endoplasmic reticulum stress in drug-induced toxicity. Pharmacology Research and Perspectives, 4(1), e00211. https://doi.org/10.1002/prp2.211

Fredriksson, L., Wink, S., Herpers, B., Benedetti, G., Hadi, M., De Bont, H., Groothuis, G., Luijten, M., Danen, E., De Graauw, M., Meerman, J., & van de Water, B. (2014). Drug-induced endoplasmic reticulum and oxidative stress responses independently sensitize toward TNFα-mediated hepatotoxicity. Toxicological Sciences, 140(1), 144–159. https://doi.org/10.1093/toxsci/kfu072

Garcia-Alonso, L., Holland, C. H., Ibrahim, M. M., Turei, D., & Saez-Rodriguez, J. (2019). Benchmark and integration of resources for the estimation of human transcription factor activities. Genome Research, 29(8), 1363–1375. https://doi.org/10.1101/gr.240663.118

Garcia-Alonso, L., Iorio, F., Matchan, A., Fonseca, N., Jaaks, P., Peat, G., Pignatelli, M., Falcone, F., Benes, C. H., Dunham, I., Bignell, G., McDade, S. S., Garnett, M. J., & Saez-Rodriguez, J. (2018). Transcription factor activities enhance markers of drug sensitivity in cancer. Cancer Research, 78(3), 769–780. https://doi.org/10.1158/0008-5472.CAN-17-1679

García-Ruiz, C., & Fernández-Checa, J. C. (2018). Mitochondrial Oxidative Stress and Antioxidants Balance in Fatty Liver Disease. Hepatology Communications, 2(12), 1425–1439. https://doi.org/10.1002/hep4.1271

Hetz, C., Zhang, K., & Kaufman, R. J. (2020). Mechanisms, regulation and functions of the unfolded protein response. In Nature Reviews Molecular Cell Biology (Vol. 21, Issue 8, pp. 421–438). Nature Research. https://doi.org/10.1038/s41580-020-0250-z

Ideker, T., Dutkowski, J., & Hood, L. (2011). Boosting signal-to-noise in complex biology: Prior knowledge is power. In Cell (Vol. 144, Issue 6, pp. 860–863). Elsevier. https://doi.org/10.1016/j.cell.2011.03.007

Igarashi, Y., Nakatsu, N., Yamashita, T., Ono, A., Ohno, Y., Urushidani, T., & Yamada, H. (2015). Open TG-GATEs: A large-scale toxicogenomics database. Nucleic Acids Research, 43(D1), D921–D927. https://doi.org/10.1093/nar/gku955

Kamburov, A., Stelzl, U., Lehrach, H., & Herwig, R. (2013). The ConsensusPathDB interaction database: 2013 Update. Nucleic Acids Research, 41(D1). https://doi.org/10.1093/nar/gks1055

Karin, M., & Dhar, D. (2016). Liver carcinogenesis: From naughty chemicals to soothing fat and the surprising role of NRF2. Carcinogenesis, 37(6), 541–546. https://doi.org/10.1093/carcin/bgw060

Kawamoto, T., Ito, Y., Morita, O., & Honda, H. (2017). Mechanism-based risk assessment strategy for drug-induced cholestasis using the transcriptional benchmark dose derived by toxicogenomics. Journal of Toxicological Sciences, 42(4), 427–436. https://doi.org/10.2131/jts.42.427

Koido, M., Kawakami, E., Fukumura, J., Noguchi, Y., Ohori, M., Nio, Y., Nicoletti, P., Aithal, G. P., Daly, A. K., Watkins, P. B., Anayama, H., Dragan, Y., Shinozawa, T., & Takebe, T. (2020). Polygenic architecture informs potential vulnerability to drug-induced liver injury. Nature Medicine, 26(10), 1541–1548. https://doi.org/10.1038/s41591-020-1023-0

Kolde, R., & Kolde, M. R. (2015). Package ‘pheatmap.’ R Package, 1(7).

Krewski, D., Andersen, M. E., Tyshenko, M. G., Krishnan, K., Hartung, T., Boekelheide, K., Wambaugh, J. F., Jones, D., Whelan, M., Thomas, R., Yauk, C., Barton-Maclaren, T., & Cote, I. (2020). Toxicity testing in the 21st century: progress in the past decade and future perspectives. Archives of Toxicology, 94(1). https://doi.org/10.1007/s00204-019-02613-4

Langfelder, P., & Horvath, S. (2008). WGCNA: An R package for weighted correlation network analysis. BMC Bioinformatics, 9(1), 559. https://doi.org/10.1186/1471-2105-9-559

Langfelder, P., Luo, R., Oldham, M. C., & Horvath, S. (2011). Is my network module preserved and reproducible? PLoS Computational Biology, 7(1), 1001057. https://doi.org/10.1371/journal.pcbi.1001057

Lanzoni, A., Castoldi, A. F., Kass, G. E. N., Terron, A., De Seze, G., Bal-Price, A., Bois, F. Y., Delclos, K. B., Doerge, D. R., Fritsche, E., Halldorsson, T., Kolossa-Gehring, M., Hougaard Bennekou, S., Koning, F., Lampen, A., Leist, M., Mantus, E., Rousselle, C., Siegrist, M., … Younes, M. (2019). Advancing human health risk assessment. EFSA Journal, 17(S1), 170712. https://doi.org/10.2903/j.efsa.2019.e170712

Liu, Z., Huang, R., Roberts, R., & Tong, W. (2019). Toxicogenomics: A 2020 Vision. In Trends in Pharmacological Sciences (Vol. 40, Issue 2, pp. 92–103). Elsevier Ltd. https://doi.org/10.1016/j.tips.2018.12.001

Love, M. I., Huber, W., & Anders, S. (2014). Moderated estimation of fold change and dispersion for RNA-seq data with DESeq2. Genome Biology, 15(12), 550. https://doi.org/10.1186/s13059-014-0550-8

MacRae, S. L., Croken, M. M. K., Calder, R. B., Aliper, A., Milholland, B., White, R. R., Zhavoronkov, A., Gladyshev, V. N., Seluanov, A., Gorbunova, V., Zhang, Z. D., & Vijg, J. (2015). DNA repair in species with extreme lifespan differences. Aging, 7(12), 1171–1184. https://doi.org/10.18632/aging.100866

Mandrekar, P., Catalano, D., Jeliazkova, V., & Kodys, K. (2008). Alcohol exposure regulates heat shock transcription factor binding and heat shock proteins 70 and 90 in monocytes and macrophages: implication for TNF-α regulation. Journal of Leukocyte Biology, 84(5), 1335–1345. https://doi.org/10.1189/jlb.0407256

Martin, L. J., & Chang, Q. (2018). DNA damage response and repair, DNA methylation, and cell death in human neurons and experimental animal neurons are different. Journal of Neuropathology and Experimental Neurology, 77(7), 636–655. https://doi.org/10.1093/jnen/nly040

Mav, D., Phadke, D. P., Balik-Meisner, M. R., Merrick, B. A., Auerbach, S., Niemeijer, M., Huppelschoten, S., Baze, A., Parmentier, C., Richert, L., van de Water, B., Shah, R. R., & Paules, R. S. (2020). Utility of Extrapolating Human S1500+ Genes to the Whole Transcriptome: Tunicamycin Case Study. Bioinformatics and Biology Insights, 14, 117793222095274. https://doi.org/10.1177/1177932220952742

Mav, D., Shah, R. R., Howard, B. E., Auerbach, S. S., Bushel, P. R., Collins, J. B., Gerhold, D. L., Judson, R. S., Karmaus, A. L., Maull, E. A., Mendrick, D. L., Merrick, B. A., Sipes, N. S., Svoboda, D., & Paules, R. S. (2018). A hybrid gene selection approach to create the S1500+ targeted gene sets for use in high-throughput transcriptomics. PLoS ONE, 13(2). https://doi.org/10.1371/journal.pone.0191105

Monroe, J. J., Tanis, K. Q., Podtelezhnikov, A. A., Nguyen, T., Machotka, S. V., Lynch, D., Evers, R., Palamanda, J., Miller, R. R., Pippert, T., Cabalu, T. D., Johnson, T. E., Aslamkhan, A. G., Kang, W., Tamburino, A. M., Mitra, K., Agrawal, N. G. B., & Sistare, F. D. (2020). Application of a Rat Liver Drug Bioactivation Transcriptional Response Assay Early in Drug Development That Informs Chemically Reactive Metabolite Formation and Potential for Drug-induced Liver Injury. Toxicological Sciences, 177(1), 281–299. https://doi.org/10.1093/toxsci/kfaa088

Morin, M. J., & Bernacki, R. J. (1983). Biochemical effects and therapeutic potential of tunicamycin in murine L1210 leukemia. Cancer Research, 43(4), 1669–1674.

Onakpoya, I. J., Heneghan, C. J., & Aronson, J. K. (2016). Post-marketing withdrawal of 462 medicinal products because of adverse drug reactions: A systematic review of the world literature. BMC Medicine, 14(1), 10. https://doi.org/10.1186/s12916-016-0553-2

Osataphan, N., Phrommintikul, A., Chattipakorn, S. C., & Chattipakorn, N. (2020). Effects of doxorubicin-induced cardiotoxicity on cardiac mitochondrial dynamics and mitochondrial function: Insights for future interventions. Journal of Cellular and Molecular Medicine, 24(12), 6534–6557. https://doi.org/10.1111/jcmm.15305

Pakos-Zebrucka, K., Koryga, I., Mnich, K., Ljujic, M., Samali, A., & Gorman, A. M. (2016). The integrated stress response. EMBO Reports, 17(10), 1374–1395. https://doi.org/10.15252/embr.201642195

Paradis, E., & Schliep, K. (2019). Ape 5.0: An environment for modern phylogenetics and evolutionary analyses in R. Bioinformatics, 35(3), 526–528. https://doi.org/10.1093/bioinformatics/bty633

Parish, S. T., Aschner, M., Casey, W., Corvaro, M., Embry, M. R., Fitzpatrick, S., Kidd, D., Kleinstreuer, N. C., Lima, B. S., Settivari, R. S., Wolf, D. C., Yamazaki, D., & Boobis, A. (2020). An evaluation framework for new approach methodologies (NAMs) for human health safety assessment. Regulatory Toxicology and Pharmacology, 112, 104592. https://doi.org/10.1016/j.yrtph.2020.104592

Peng, C., Stewart, A. G., Woodman, O. L., Ritchie, R. H., & Qin, C. X. (2020). Non-Alcoholic Steatohepatitis: A Review of Its Mechanism, Models and Medical Treatments. In Frontiers in Pharmacology (Vol. 11). Frontiers Media S.A. https://doi.org/10.3389/fphar.2020.603926

Perkins, E., Garcia-Reyero, N., Edwards, S., Wittwehr, C., Villeneuve, D., Lyons, D., & Ankley, G. (2015). The adverse outcome pathway: A conceptual framework to support toxicity testing in the twenty-first century. In Computational Systems Toxicology (pp. 1–26). Springer New York. https://doi.org/10.1007/978-1-4939-2778-4_1

Peter Langfelder, A., Hor-, S., Cai, C., Dong, J., Miller, J., Song, L., Yip, A., & Zhang Maintainer Peter Langfelder, B. (2020). Package “WGCNA” Title Weighted Correlation Network Analysis. http:

Phillips, J. R., Svoboda, D. L., Tandon, A., Patel, S., Sedykh, A., Mav, D., Kuo, B., Yauk, C. L., Yang, L., Thomas, R. S., Gift, J. S., Allen Davis, J., Olszyk, L., Alex Merrick, B., Paules, R. S., Parham, F., Saddler, T., Shah, R. R., & Auerbach, S. S. (2019). BMD Express 2: Enhanced transcriptomic dose-response analysis workflow. Bioinformatics, 35(10), 1780–1782. https://doi.org/10.1093/bioinformatics/bty878

Podtelezhnikov, A. A., Monroe, J. J., Aslamkhan, A. G., Pearson, K., Qin, C., Tamburino, A. M., Loboda, A. P., Glaab, W. E., Sistare, F. D., & Tanis, K. Q. (2020). Quantitative Transcriptional Biomarkers of Xenobiotic Receptor Activation in Rat Liver for the Early Assessment of Drug Safety Liabilities. Toxicological Sciences, 175(1), 98–112. https://doi.org/10.1093/toxsci/kfaa026

Rana, P., Aleo, M. D., Gosink, M., & Will, Y. (2019). Evaluation of in Vitro Mitochondrial Toxicity Assays and Physicochemical Properties for Prediction of Organ Toxicity Using 228 Pharmaceutical Drugs. In Chemical Research in Toxicology (Vol. 32, Issue 1, pp. 156–167). American Chemical Society. https://doi.org/10.1021/acs.chemrestox.8b00246

Reuben, A., Tillman, H., Fontana, R. J., Davern, T., Mcguire, B., Stravitz, R. T., Durkalski, V., Larson, A. M., Liou, I., Fix, O., Schilsky, M., Mccashland, T., Hay, J. E., Murray, N., Shaikh, O. S., Ganger, D., Zaman, A., Han, S. B., Chung, R. T., … Lee, W. M. (2016). Outcomes in adults with acute liver failure between 1998 and 2013: An observational cohort study. Annals of Internal Medicine, 164(11), 724–732. https://doi.org/10.7326/M15-2211

Rezzani, R. (2004). Cyclosporine A and adverse effects on organs: Histochemical studies. Progress in Histochemistry and Cytochemistry, 39(2), 85–128. https://doi.org/10.1016/j.proghi.2004.04.001

Rivetti, C., Allen, T. E. H., Brown, J. B., Butler, E., Carmichael, P. L., Colbourne, J. K., Dent, M., Falciani, F., Gunnarsson, L., Gutsell, S., Harrill, J. A., Hodges, G., Jennings, P., Judson, R., Kienzler, A., Margiotta-Casaluci, L., Muller, I., Owen, S. F., Rendal, C., … Campos, B. (2020). Vision of a near future: Bridging the human health–environment divide. Toward an integrated strategy to understand mechanisms across species for chemical safety assessment. In Toxicology in Vitro (Vol. 62). Elsevier Ltd. https://doi.org/10.1016/j.tiv.2019.104692

Ron, D., & Walter, P. (2007). Signal integration in the endoplasmic reticulum unfolded protein response. In Nature Reviews Molecular Cell Biology (Vol. 8, Issue 7, pp. 519–529). Nature Publishing Group. https://doi.org/10.1038/nrm2199

RStudio Inc. (2014). shiny: Web Application Framework for R. R package version 0.9.1. In http://CRAN.R-project.org/package=shiny.

Safe, S., Jin, U. H., Park, H., Chapkin, R. S., & Jayaraman, A. (2020). Aryl hydrocarbon receptor (AHR) ligands as selective ahr modulators (SAHRMS). In International Journal of Molecular Sciences (Vol. 21, Issue 18, pp. 1–16). MDPI AG. https://doi.org/10.3390/ijms21186654

Sax, N. I. (1975). Dangerous properties of industrial materials. In Van Nostrand Reinhold: Vol. £21.25 (12th ed.). John Wiley & Sons, Ltd. https://doi.org/10.2105/ajph.54.5.866-b

Shannon, P., Markiel, A., Ozier, O., Baliga, N. S., Wang, J. T., Ramage, D., Amin, N., Schwikowski, B., & Ideker, T. (2003). Cytoscape: A software Environment for integrated models of biomolecular interaction networks. Genome Research, 13(11), 2498–2504. https://doi.org/10.1101/gr.1239303

Smith, J. R., Hayman, G. T., Wang, S. J., Laulederkind, S. J. F., Hoffman, M. J., Kaldunski, M. L., Tutaj, M., Thota, J., Nalabolu, H. S., Ellanki, S. L. R., Tutaj, M. A., De Pons, J. L., Kwitek, A. E., Dwinell, M. R., & Shimoyama, M. E. (2020). The Year of the Rat: The Rat Genome Database at 20: A multi-species knowledgebase and analysis platform. Nucleic Acids Research, 48(D1), D731–D742. https://doi.org/10.1093/nar/gkz1041

Solotke, M. T., Dhruva, S. S., Downing, N. S., Shah, N. D., & Ross, J. S. (2018). New and incremental FDA black box warnings from 2008 to 2015. Expert Opinion on Drug Safety, 17(2), 117–123. https://doi.org/10.1080/14740338.2018.1415323

Soufan, O., Ewald, J., Viau, C., Crump, D., Hecker, M., Basu, N., & Xia, J. (2019). T1000: A reduced gene set prioritized for toxicogenomic studies. PeerJ, 2019(10). https://doi.org/10.7717/peerj.7975

Stacklies, W., Redestig, H., Scholz, M., Walther, D., & Selbig, J. (2007). pcaMethods - A bioconductor package providing PCA methods for incomplete data. Bioinformatics, 23(9), 1164–1167. https://doi.org/10.1093/bioinformatics/btm069

Sutherland, J. J., Webster, Y. W., Willy, J. A., Searfoss, G. H., Goldstein, K. M., Irizarry, A. R., Hall, D. G., & Stevens, J. L. (2018). Toxicogenomic module associations with pathogenesis: A network-based approach to understanding drug toxicity. Pharmacogenomics Journal, 18(3), 377–390. https://doi.org/10.1038/tpj.2017.17

Sutherland, Jeffrey J., Jolly, R. A., Goldstein, K. M., & Stevens, J. L. (2016). Assessing Concordance of Drug-Induced Transcriptional Response in Rodent Liver and Cultured Hepatocytes. PLoS Computational Biology, 12(3). https://doi.org/10.1371/journal.pcbi.1004847

Szklarczyk, D., Franceschini, A., Wyder, S., Forslund, K., Heller, D., Huerta-Cepas, J., Simonovic, M., Roth, A., Santos, A., Tsafou, K. P., Kuhn, M., Bork, P., Jensen, L. J., & von Mering, C. (2015). STRING v10: protein–protein interaction networks, integrated over the tree of life. Nucleic Acids Research, 43(D1), D447–D452. https://doi.org/10.1093/nar/gku1003

Troyanskaya, O., Cantor, M., Sherlock, G., Brown, P., Hastie, T., Tibshirani, R., Botstein, D., & Altman, R. B. (2001). Missing value estimation methods for DNA microarrays. In Bioinformatics (Vol. 17, Issue 6). https://doi.org/10.1093/bioinformatics/17.6.520

Vahle, J. L., Anderson, U., Blomme, E. A. G., Hoflack, J. C., & Stiehl, D. P. (2018). Use of toxicogenomics in drug safety evaluation: Current status and an industry perspective. Regulatory Toxicology and Pharmacology, 96, 18–29. https://doi.org/10.1016/j.yrtph.2018.04.011

Van den Hof, W. F. P. M., Ruiz-Aracama, A., Van Summeren, A., Jennen, D. G. J., Gaj, S., Coonen, M. L. J., Brauers, K., Wodzig, W. K. W. H., van Delft, J. H. M., & Kleinjans, J. C. S. (2015a). Integrating multiple omics to unravel mechanisms of Cyclosporin A induced hepatotoxicity in vitro. Toxicology in Vitro, 29(3), 489–501. https://doi.org/10.1016/j.tiv.2014.12.016

Van den Hof, W. F. P. M., Ruiz-Aracama, A., Van Summeren, A., Jennen, D. G. J., Gaj, S., Coonen, M. L. J., Brauers, K., Wodzig, W. K. W. H., van Delft, J. H. M., & Kleinjans, J. C. S. (2015b). Integrating multiple omics to unravel mechanisms of Cyclosporin A induced hepatotoxicity in vitro. Toxicology in Vitro, 29(3), 489–501. https://doi.org/10.1016/j.tiv.2014.12.016

Van Summeren, A., Renes, J., Lizarraga, D., Bouwman, F. G., Noben, J. P., Van Delft, J. H. M., Kleinjans, J. C. S., & Mariman, E. C. M. (2013). Screening for drug-induced hepatotoxicity in primary mouse hepatocytes using acetaminophen, amiodarone, and cyclosporin A as model compounds: An omics-guided approach. OMICS A Journal of Integrative Biology, 17(2), 71–83. https://doi.org/10.1089/omi.2012.0079

Vickers, A. E. M., Ulyanov, A. V., & Fisher, R. L. (2017). Liver effects of clinical drugs differentiated in human liver slices. International Journal of Molecular Sciences, 18(3). https://doi.org/10.3390/ijms18030574

Watkins, P. B. (2011). Drug safety sciences and the bottleneck in drug development. Clinical Pharmacology and Therapeutics, 89(6), 788–790. https://doi.org/10.1038/clpt.2011.63

Weaver, R. J., Blomme, E. A., Chadwick, A. E., Copple, I. M., Gerets, H. H. J., Goldring, C. E., Guillouzo, A., Hewitt, P. G., Ingelman-Sundberg, M., Jensen, K. G., Juhila, S., Klingmüller, U., Labbe, G., Liguori, M. J., Lovatt, C. A., Morgan, P., Naisbitt, D. J., Pieters, R. H. H., Snoeys, J., … Park, B. K. (2020). Managing the challenge of drug-induced liver injury: a roadmap for the development and deployment of preclinical predictive models. Nature Reviews Drug Discovery, 19(2), 131–148. https://doi.org/10.1038/s41573-019-0048-x

Wickham H. (2008). Applied Spatial Data Analysis with R. In Applied Spatial Data Analysis with R. Springer New York. https://doi.org/10.1007/978-0-387-78171-6

Wink, S., Hiemstra, S., Huppelschoten, S., Danen, E., Niemeijer, M., Hendriks, G., Vrieling, H., Herpers, B., & Van De Water, B. (2014). Quantitative high content imaging of cellular adaptive stress response pathways in toxicity for chemical safety assessment. Chemical Research in Toxicology, 27(3), 338– 355. https://doi.org/10.1021/tx4004038

Wolters, J. E. J., Van Herwijnen, M. H. M., Theunissen, D. H. J., Jennen, D. G. J., Van Den Hof, W. F. P. M., De Kok, T. M. C. M., Schaap, F. G., Van Breda, S. G. J., & Kleinjans, J. C. S. (2016). Integrative “-Omics” Analysis in Primary Human Hepatocytes Unravels Persistent Mechanisms of Cyclosporine A-Induced Cholestasis. Chemical Research in Toxicology, 29(12), 2164–2174. https://doi.org/10.1021/acs.chemrestox.6b00337

Woolbright, B. (2017). The impact of sterile inflammation in acute liver injury. Journal of Clinical and Translational Research, 3(1), 170. https://doi.org/10.18053/jctres.03.2017s1.003

World Health Organization. (2017). Harmonization Project Document 11 Guidance document on evaluating and expressing uncertainty in hazard characterization. In World Health Organization. https://doi.org/ISBN 978 92 4 150761 5

Yang, Y., Nadanaciva, S., Will, Y., Woodhead, J. L., Howell, B. A., Watkins, P. B., & Siler, S. Q. (2015). MITOsym®: A mechanistic, mathematical model of hepatocellular respiration and bioenergetics. Pharmaceutical Research, 32(6), 1975–1992. https://doi.org/10.1007/s11095-014-1591-0

Yin, W., Mendoza, L., Monzon-Sandoval, J., Urrutia, A. O., & Gutierrez, H. (2021). Emergence of co-expression in gene regulatory networks. PLoS ONE, 16(4 April), e0247671. https://doi.org/10.1371/journal.pone.0247671

Yorita Christensen, K. L., Carrico, C. K., Sanyal, A. J., & Gennings, C. (2013). Multiple classes of environmental chemicals are associated with liver disease: NHANES 2003-2004. International Journal of Hygiene and Environmental Health, 216(6), 703–709. https://doi.org/10.1016/j.ijheh.2013.01.005

Zhang, B., & Horvath, S. (2005). WeightedNetwork2005.pdf. Statistical Applications in Genetics and Molecular Biology. https://pdfs.semanticscholar.org/bcaa/533fd46e4d0ba2d5de65f7bb576b6ec0a5a1.pdf

Zhang, J., & Venkat, D. (2020). Frequent Offenders and Patterns of Injury. In Clinics in Liver Disease (Vol. 24, Issue 1, pp. 37–48). W.B. Saunders. https://doi.org/10.1016/j.cld.2019.09.002

Zhang, L., Dong, Y., Wang, W., Zhao, T., Huang, T., Khan, A., Wang, L., Liu, Z., Xie, J., & Niu, B. (2020). Ethionine Suppresses Mitochondria Autophagy and Induces Apoptosis via Activation of Reactive Oxygen Species in Neural Tube Defects. Frontiers in Neurology, 11, 242. https://doi.org/10.3389/fneur.2020.00242

Zhang, S., Wang, C., Tang, S., Deng, S., Zhou, Y., Dai, C., Yang, X., & Xiao, X. (2014). Inhibition of autophagy promotes caspase-mediated apoptosis by tunicamycin in HepG2 cells. Toxicology Mechanisms and Methods, 24(9), 654–665. https://doi.org/10.3109/15376516.2014.956915

