## Supplementary figures and images for "The human hepatocyte TXG-MAPr: WGCNA transcriptomic modules to support mechanism-based risk assessment"

### Supplementary Figure S1

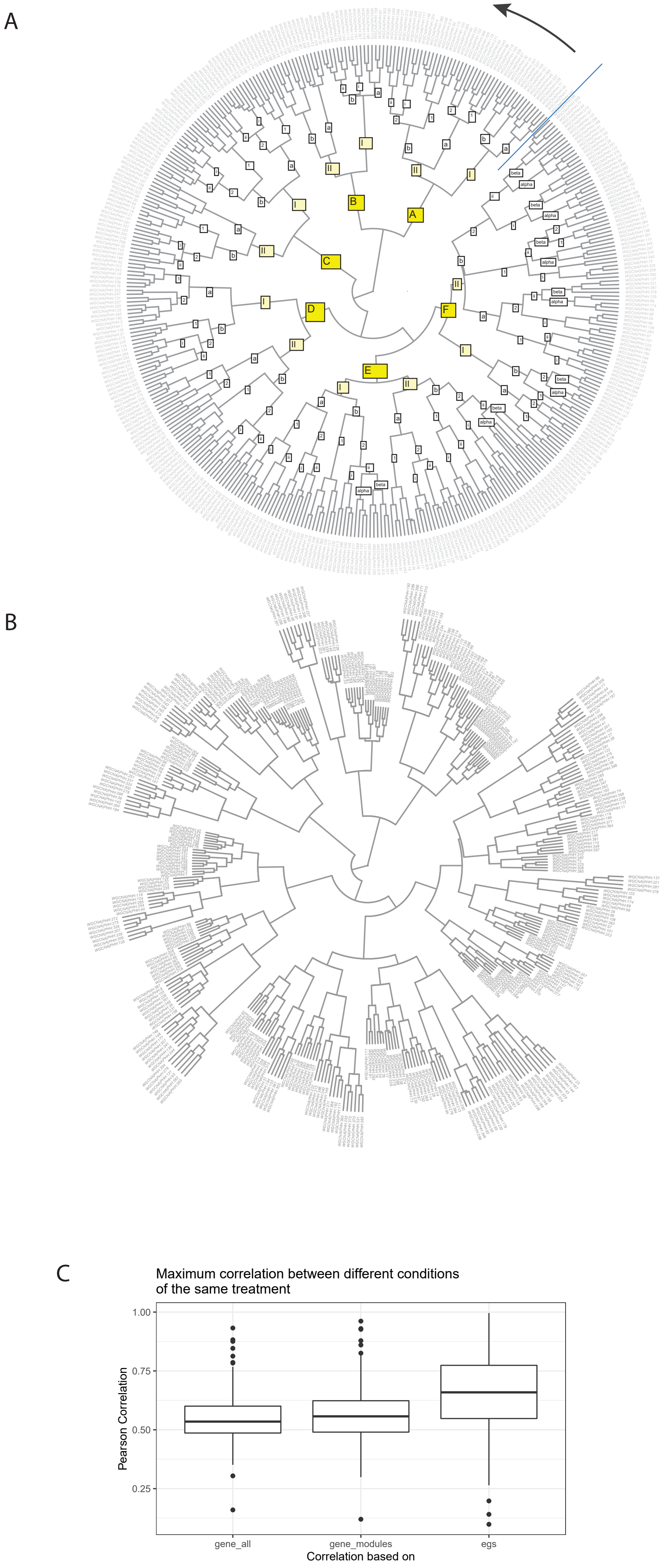

### Supplementary Figure S2

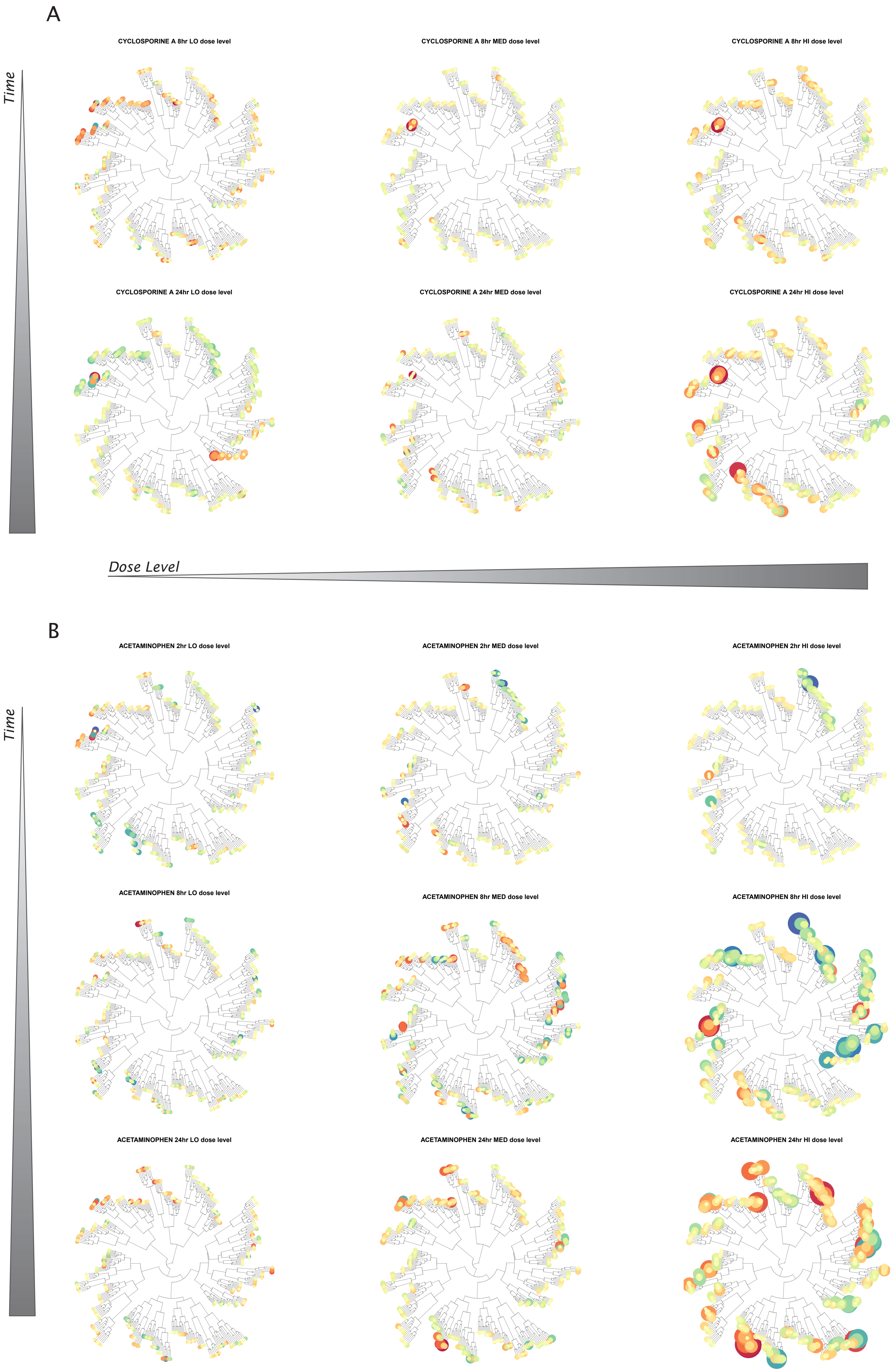

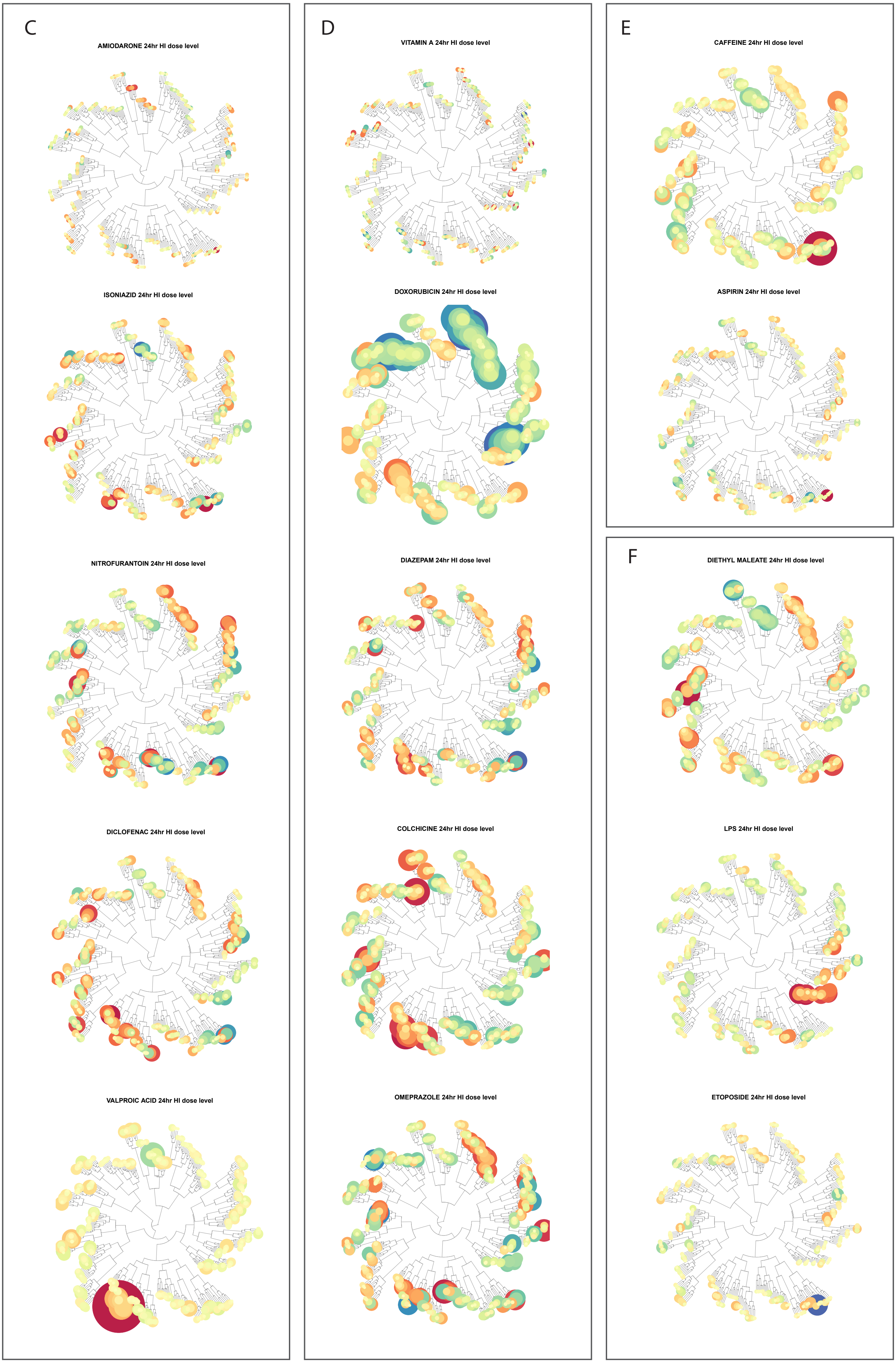

### Supplementary Figure S3

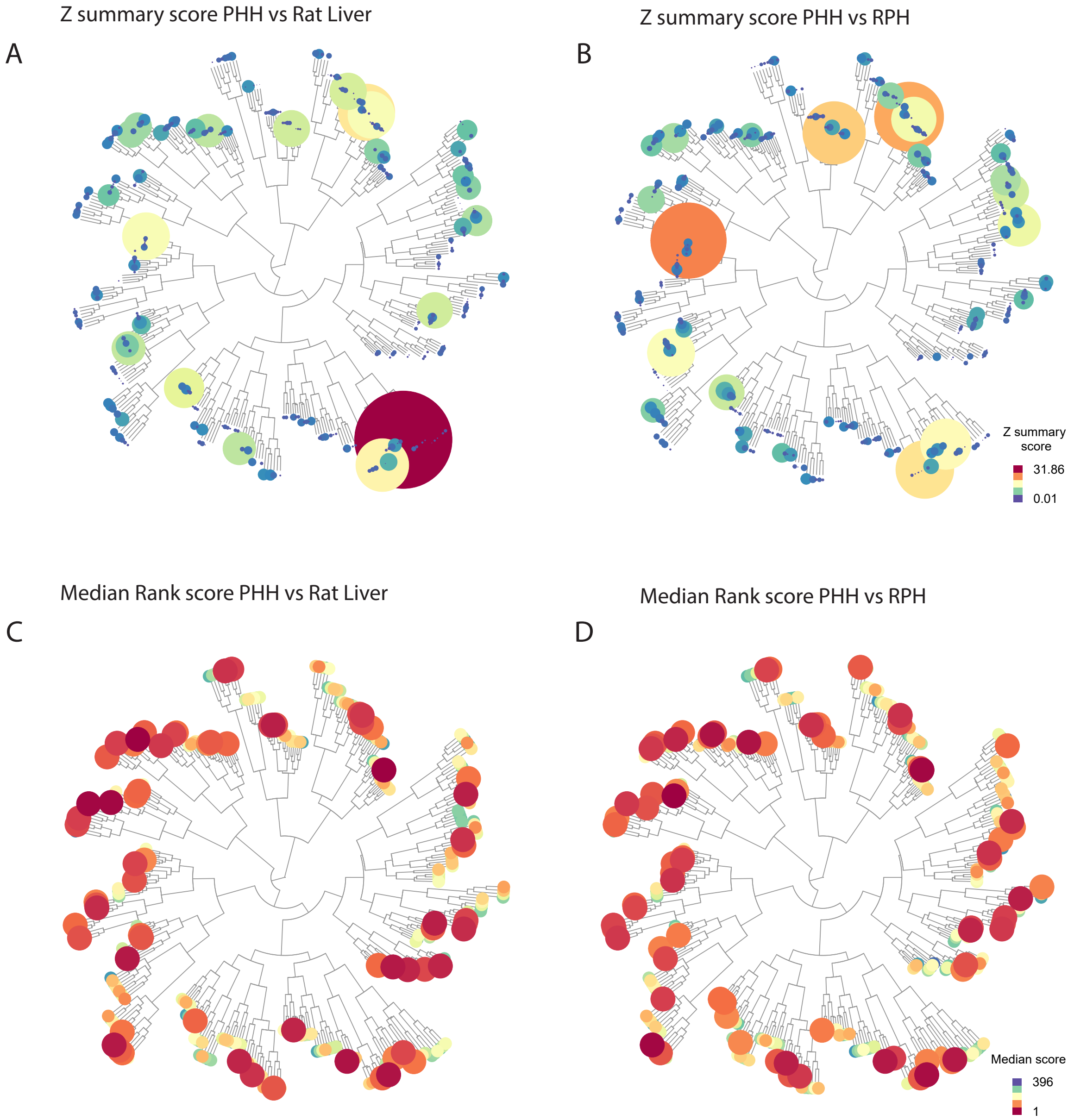

### Supplementary Figure S4

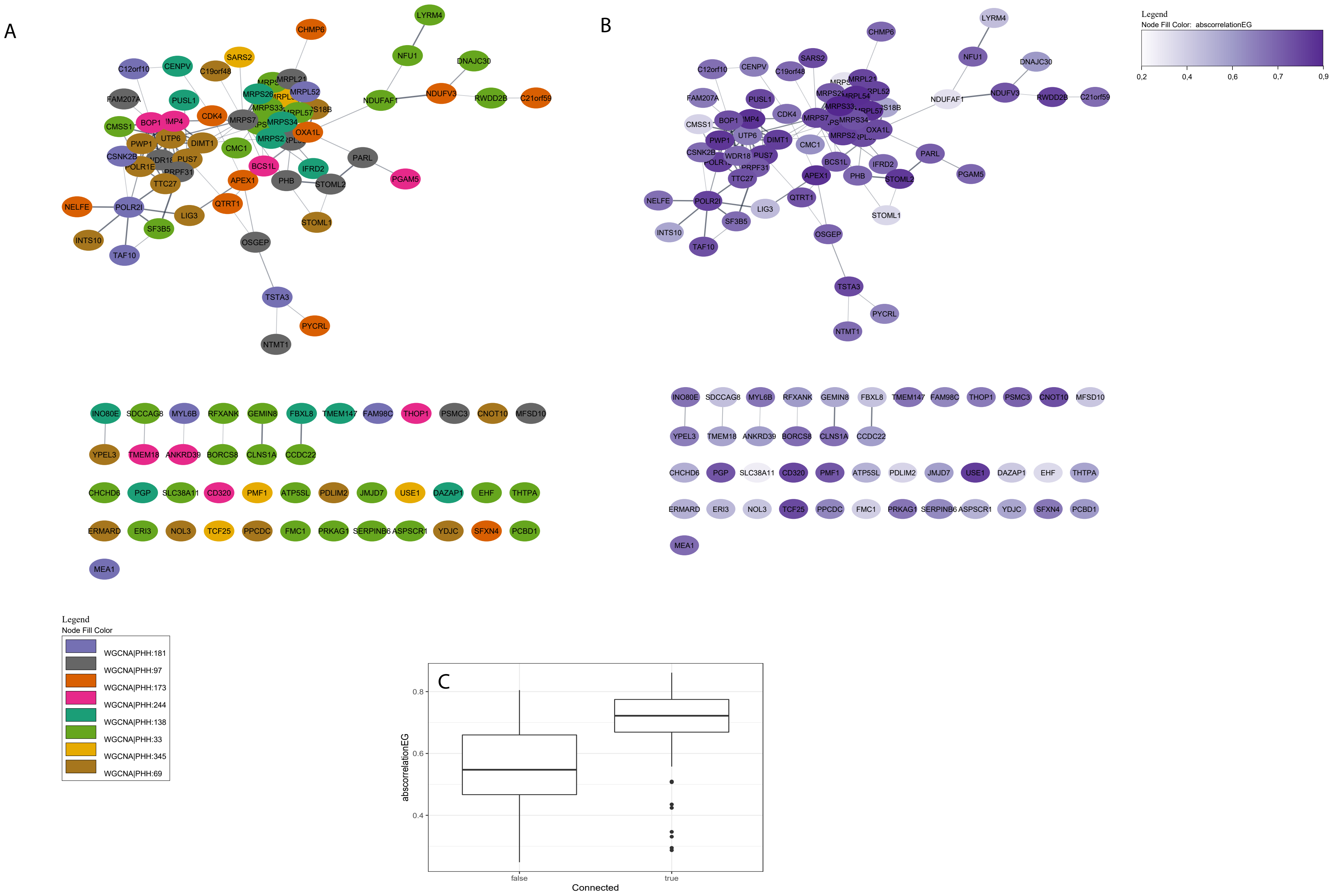

### Supplementary Figure S5

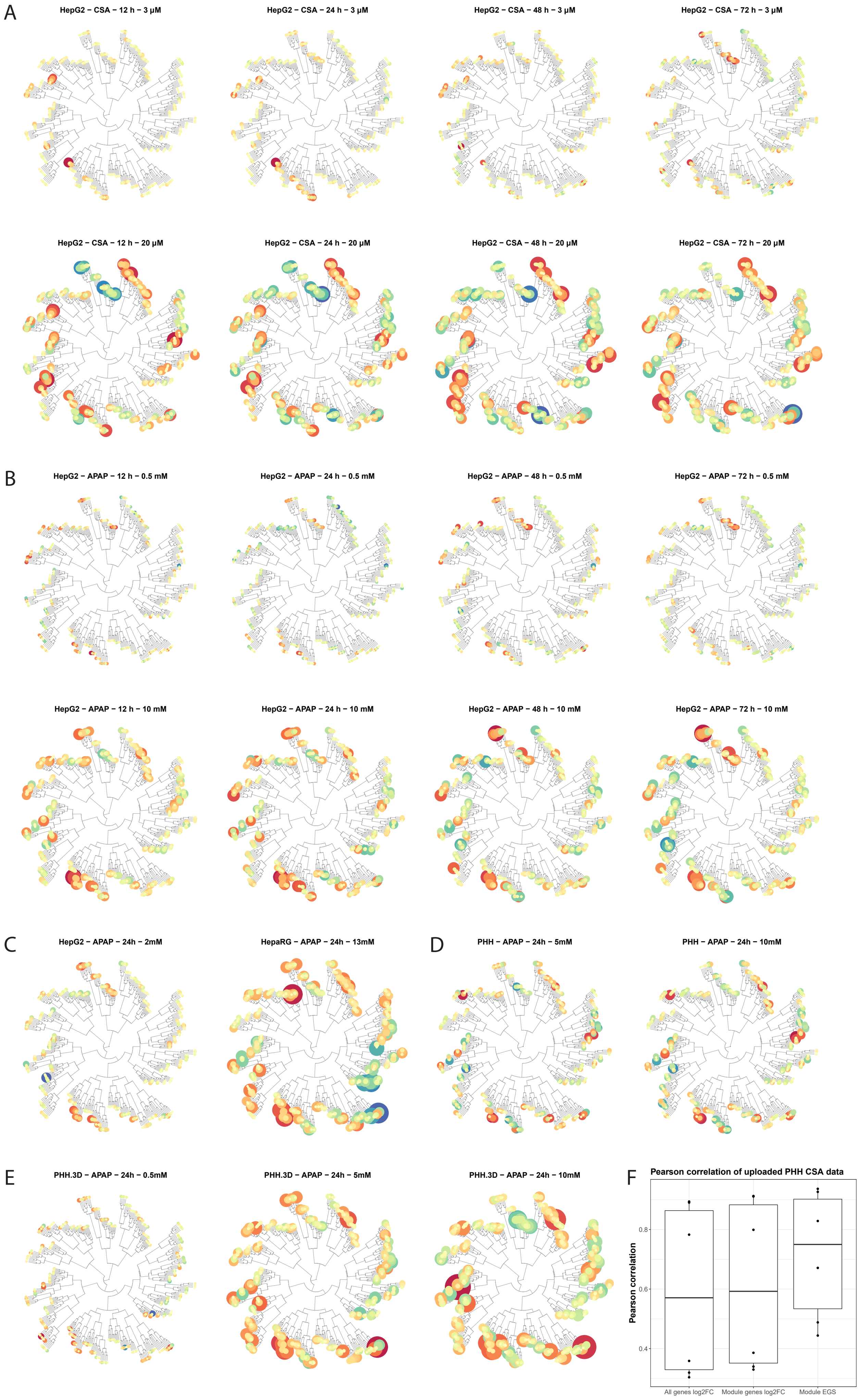

### Supplementary Figure S6

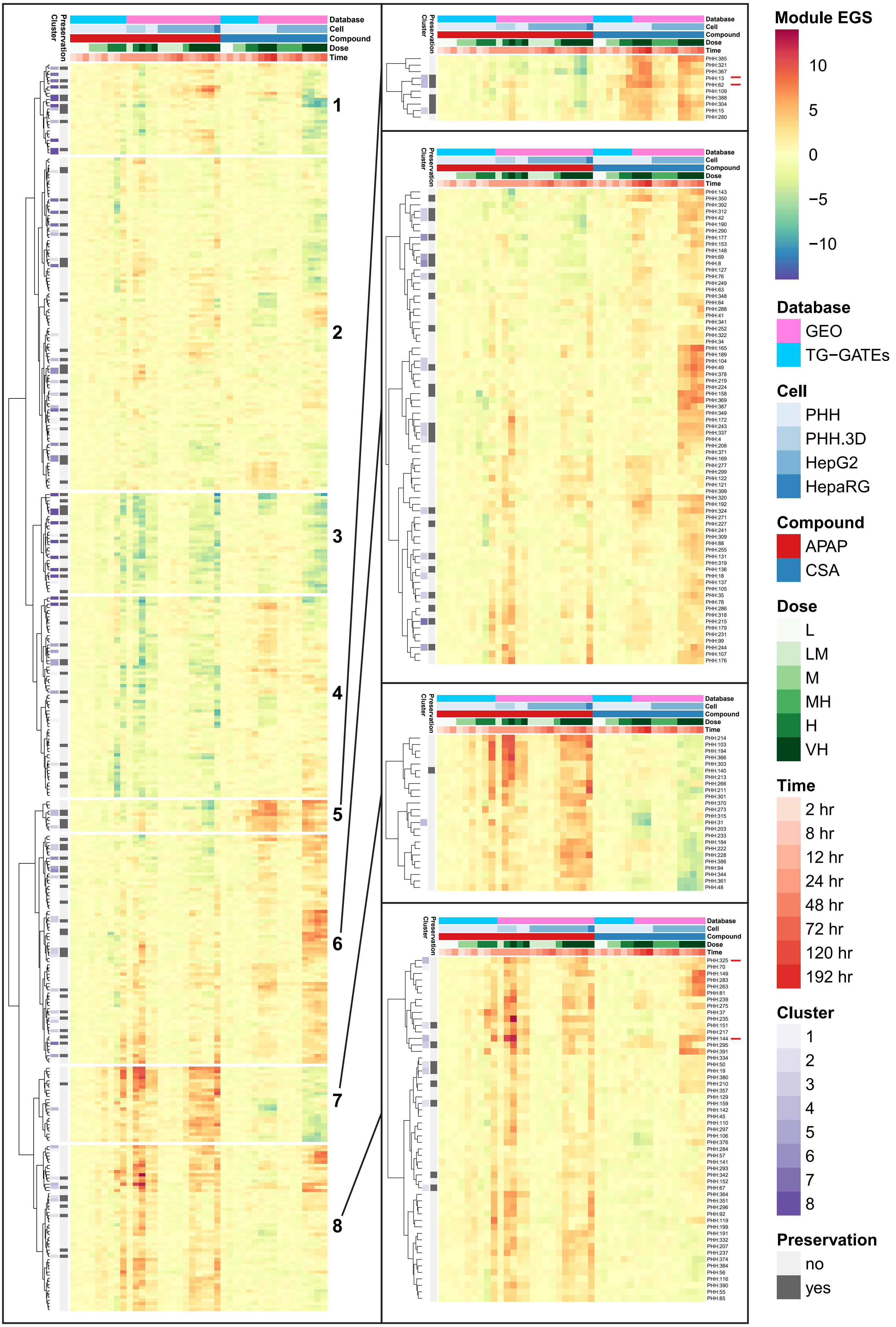

### Supplementary Figure S7

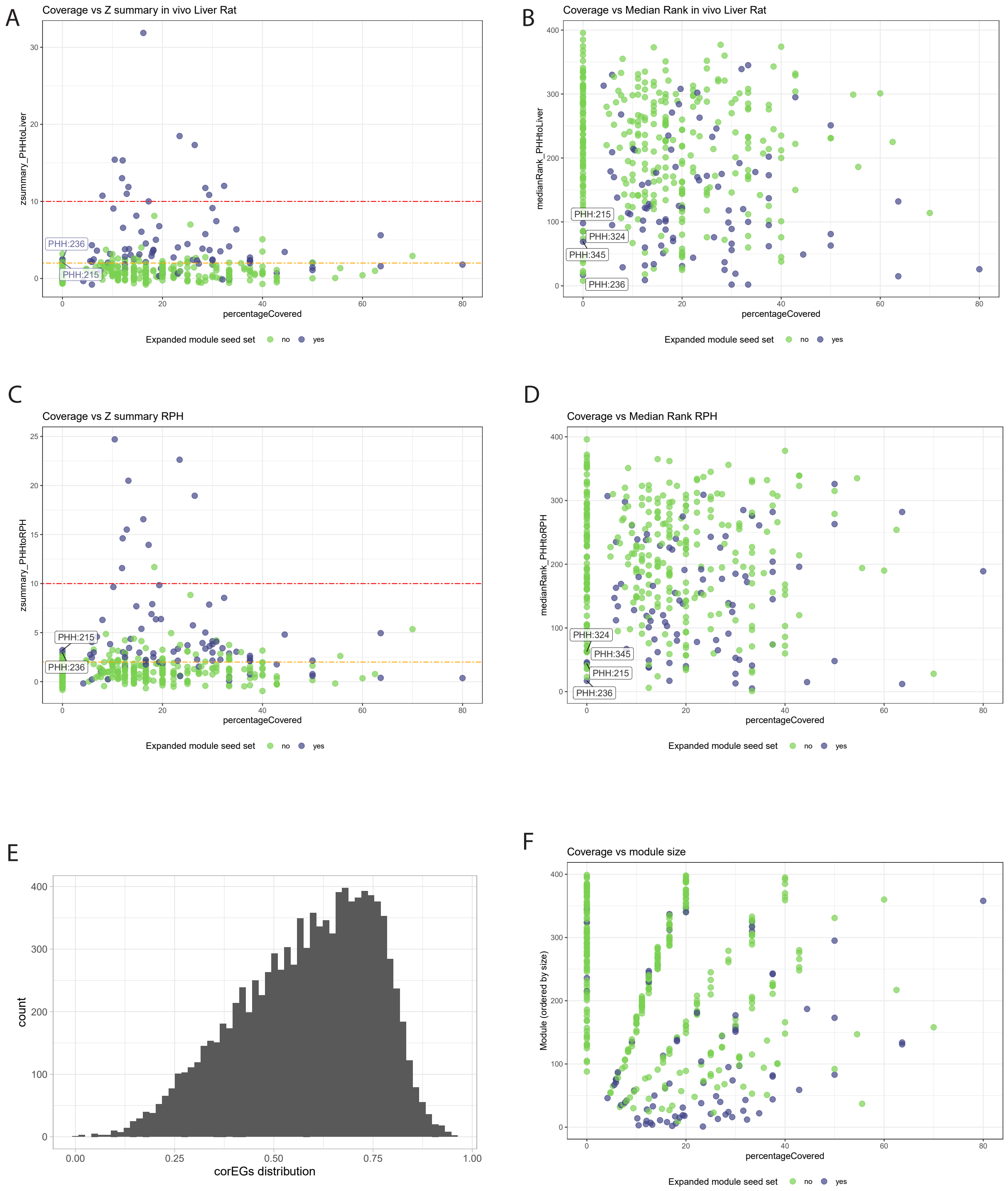

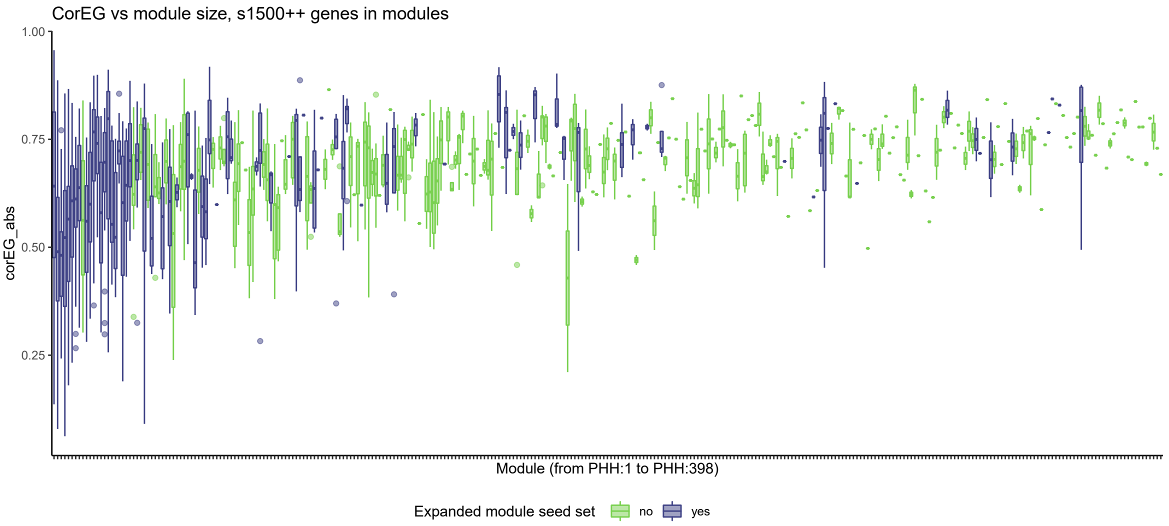

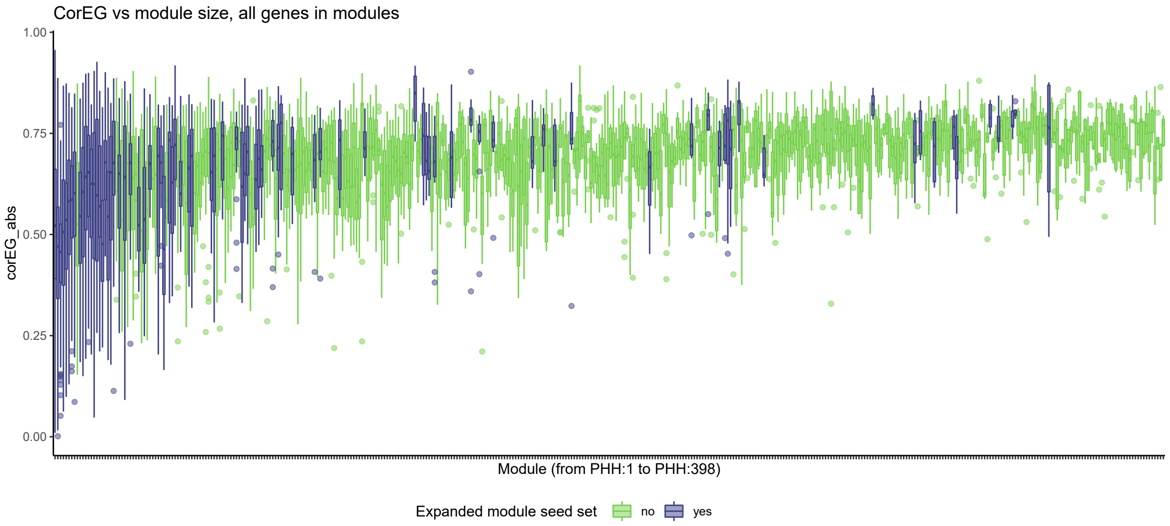

### Supplementary Figure S8

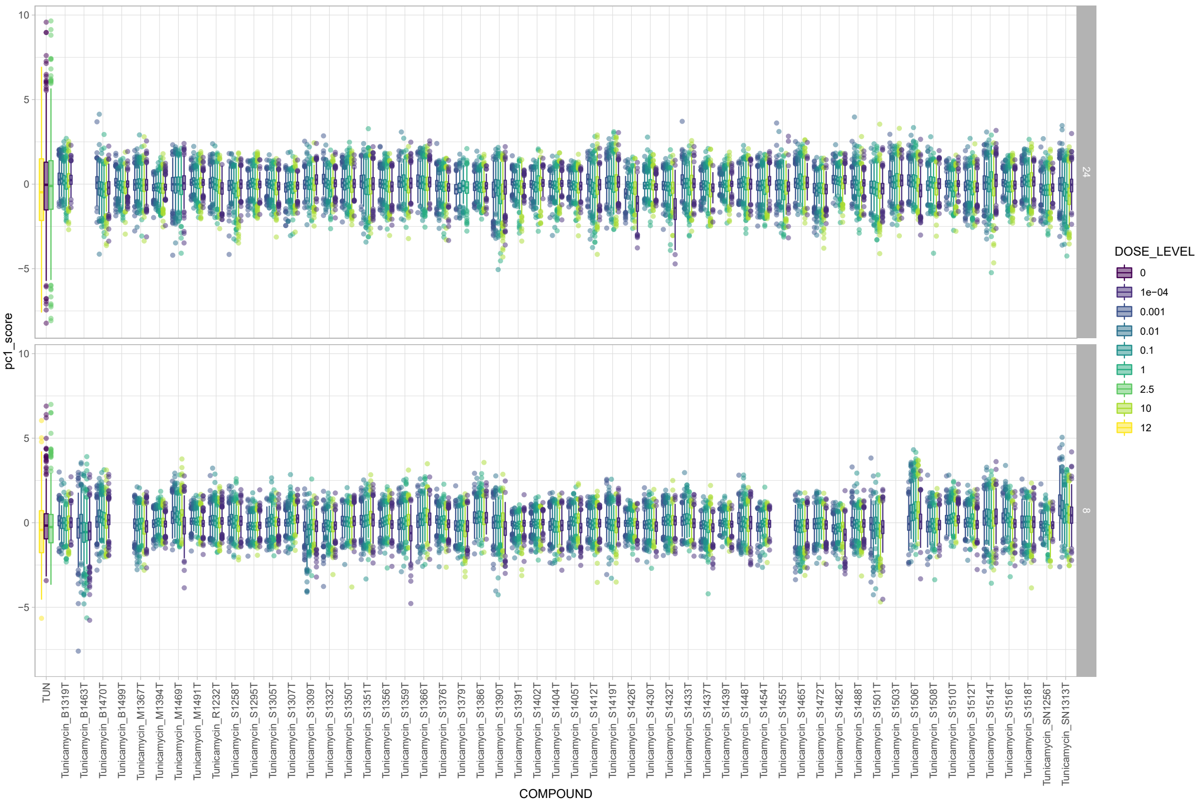

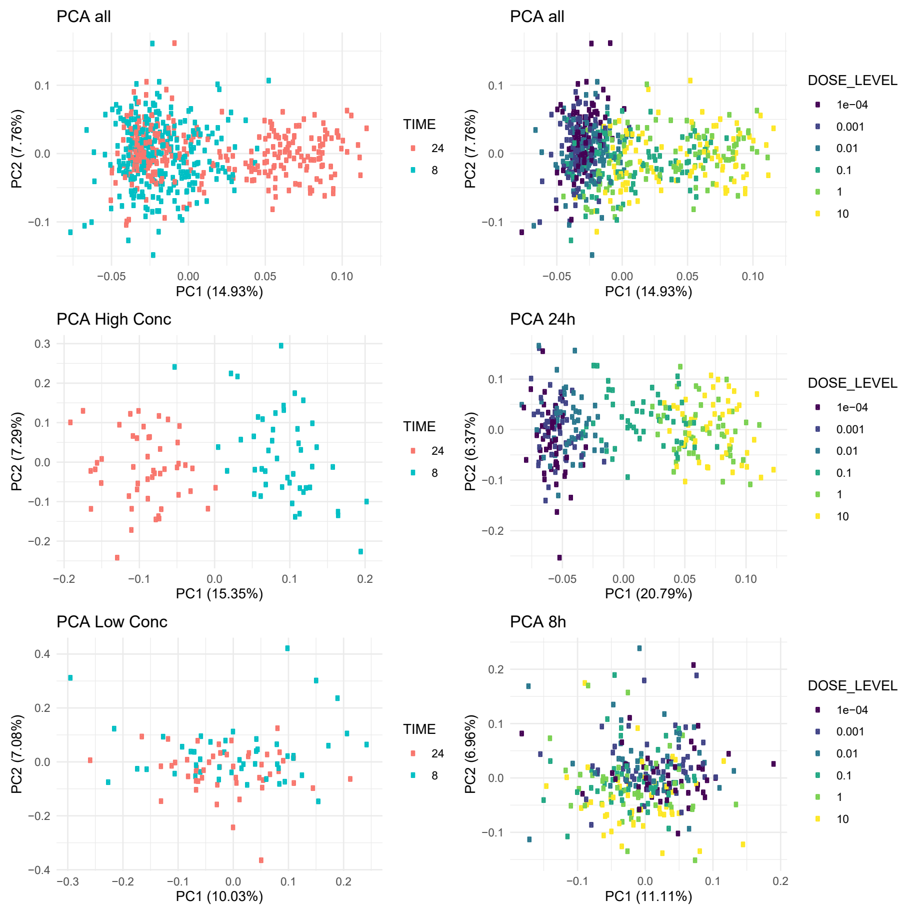

### Supplementary Figure S10

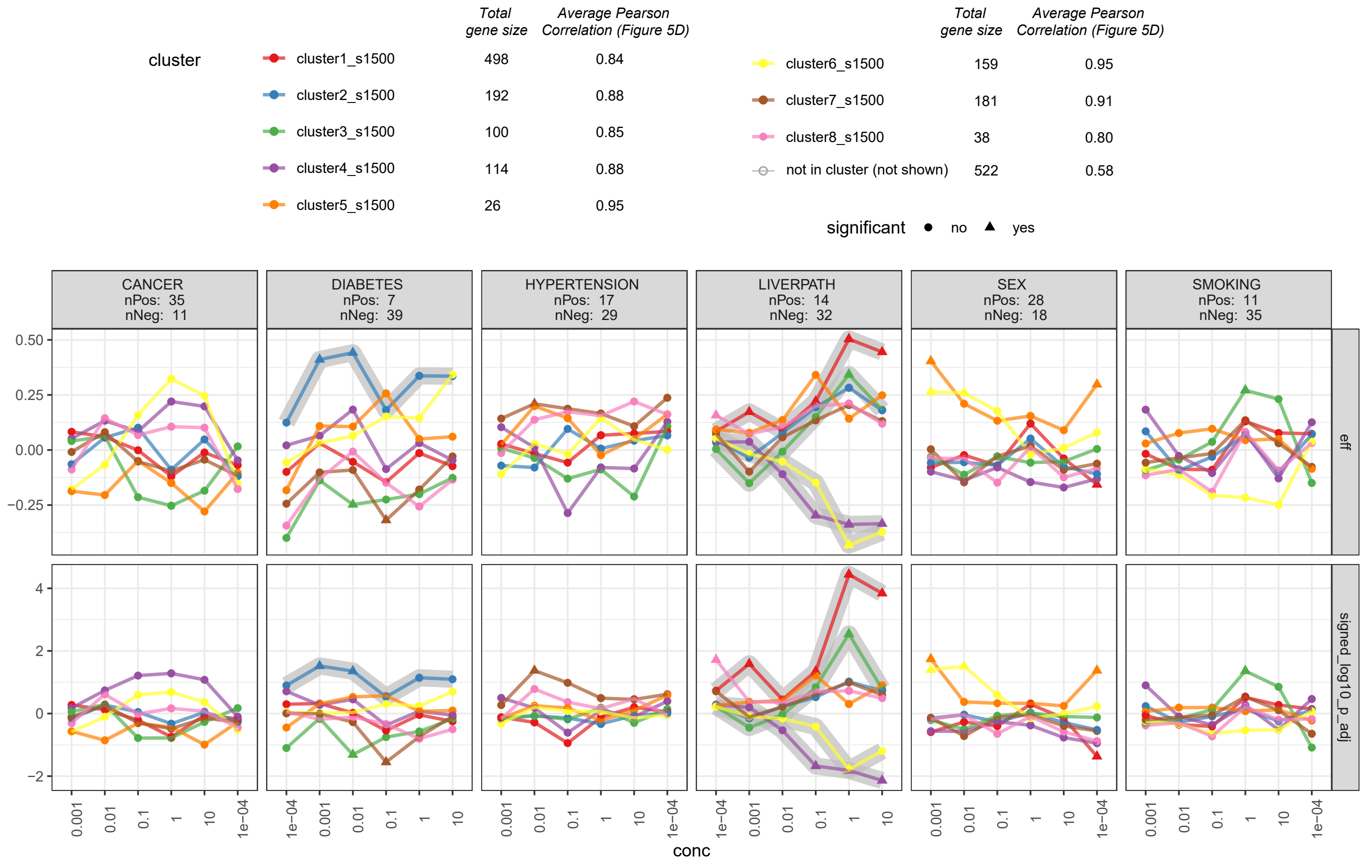

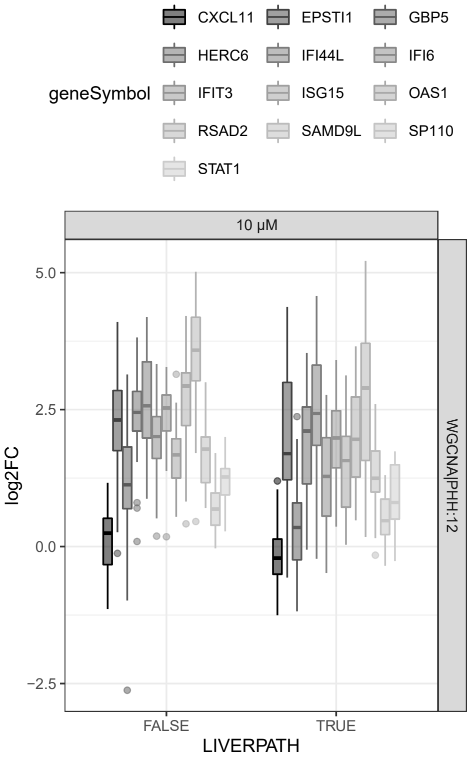

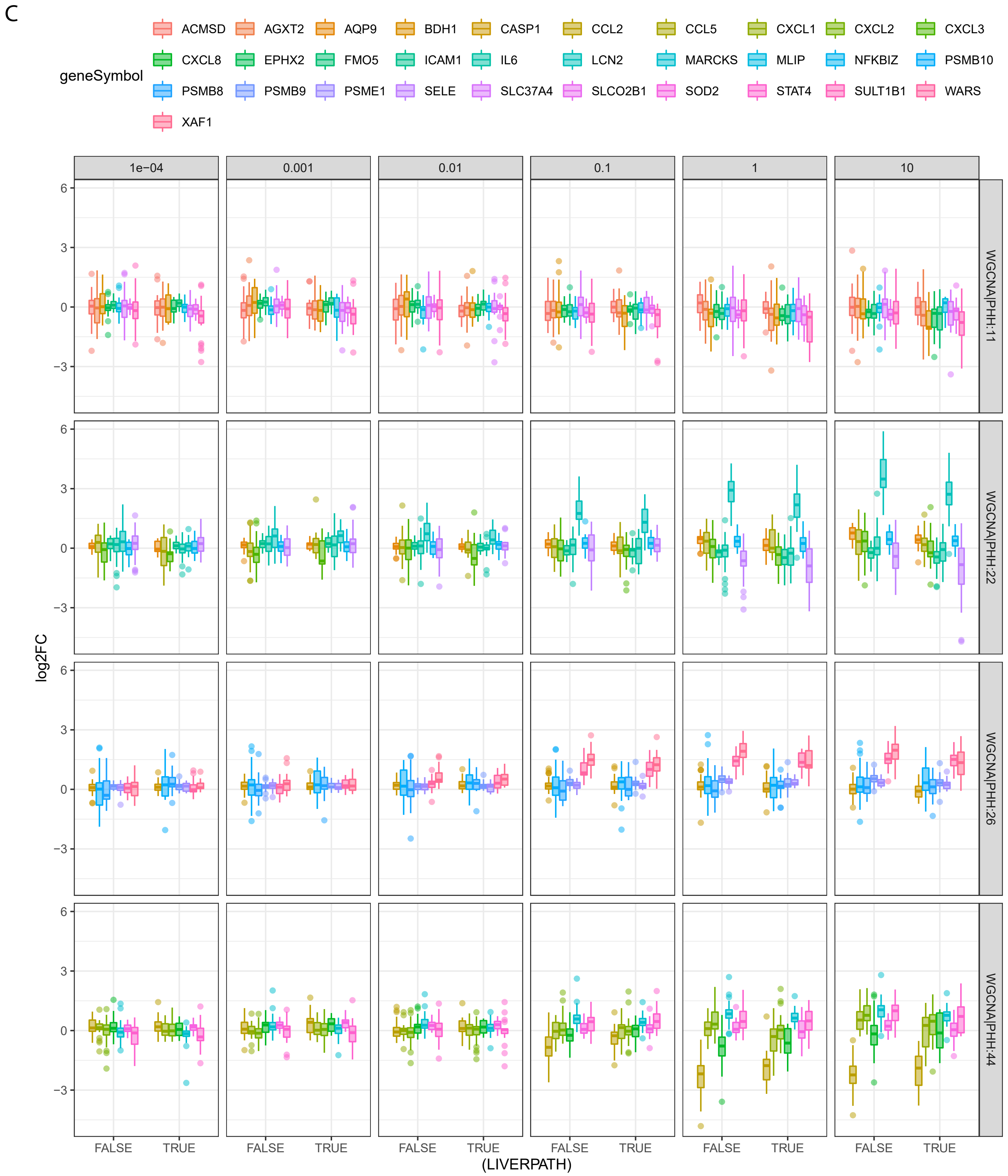

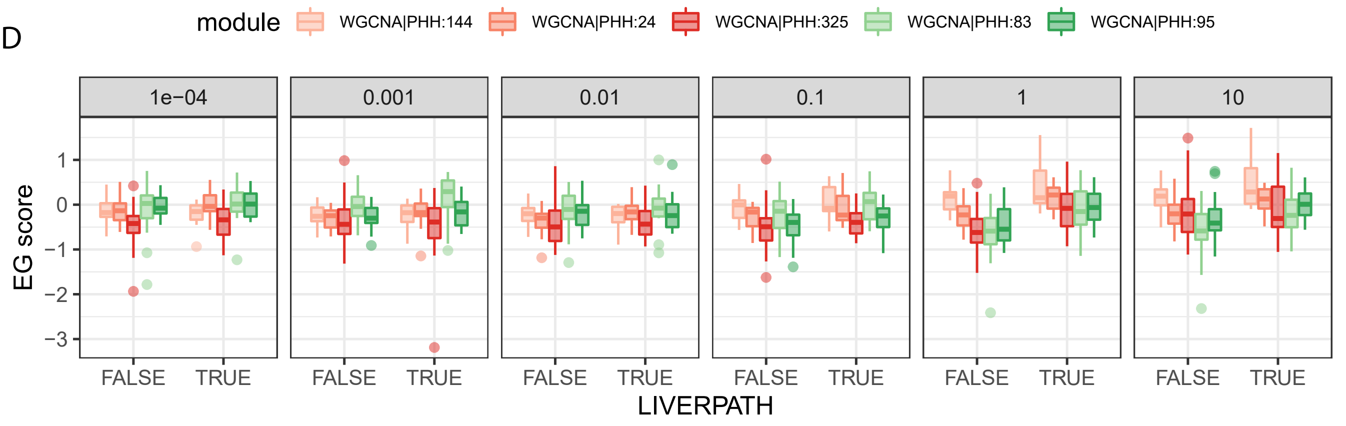

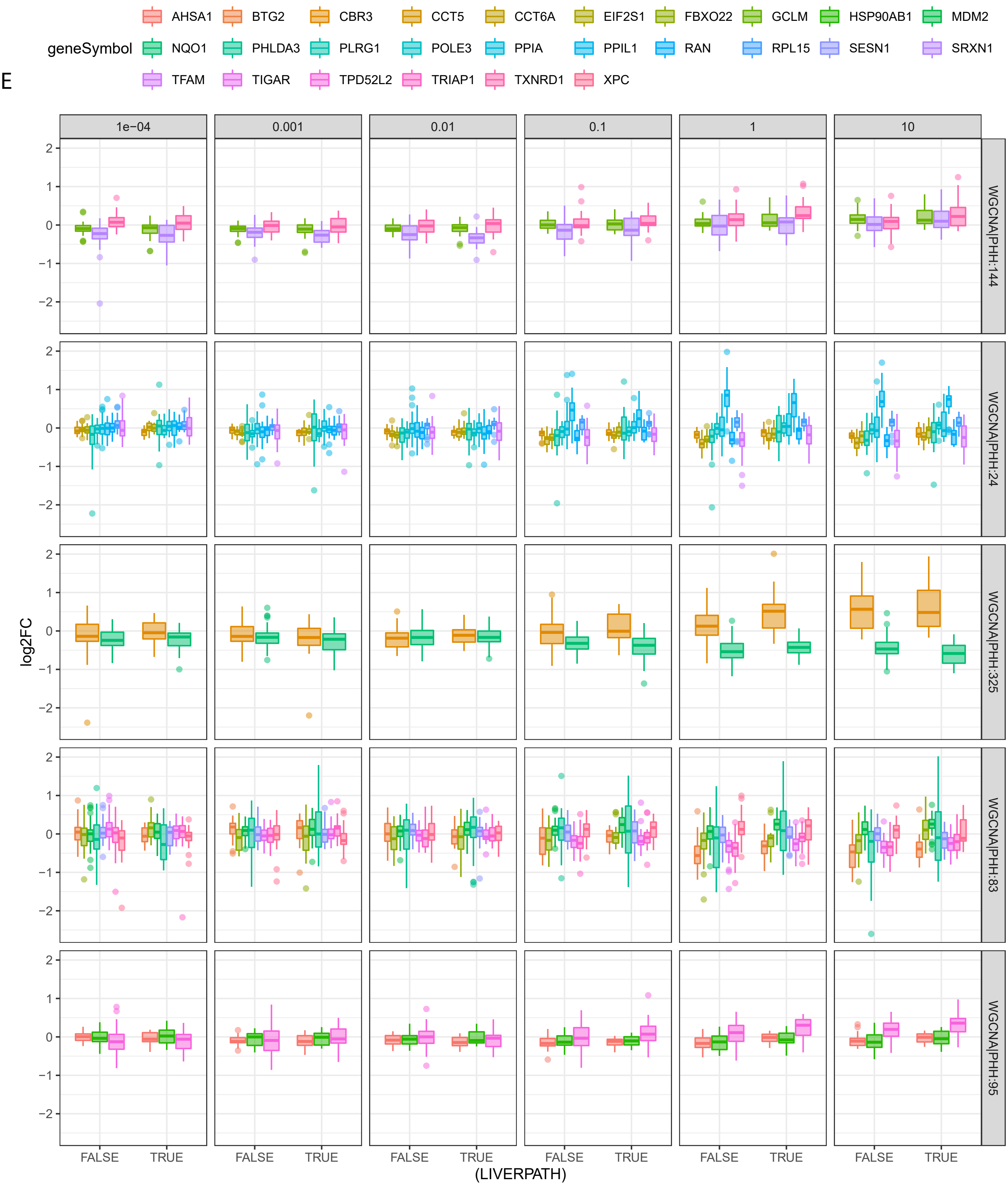
