## Supplementary Figure S9 for "The human hepatocyte TXG-MAPr: WGCNA transcriptomic modules to support mechanism-based risk assessment"

A

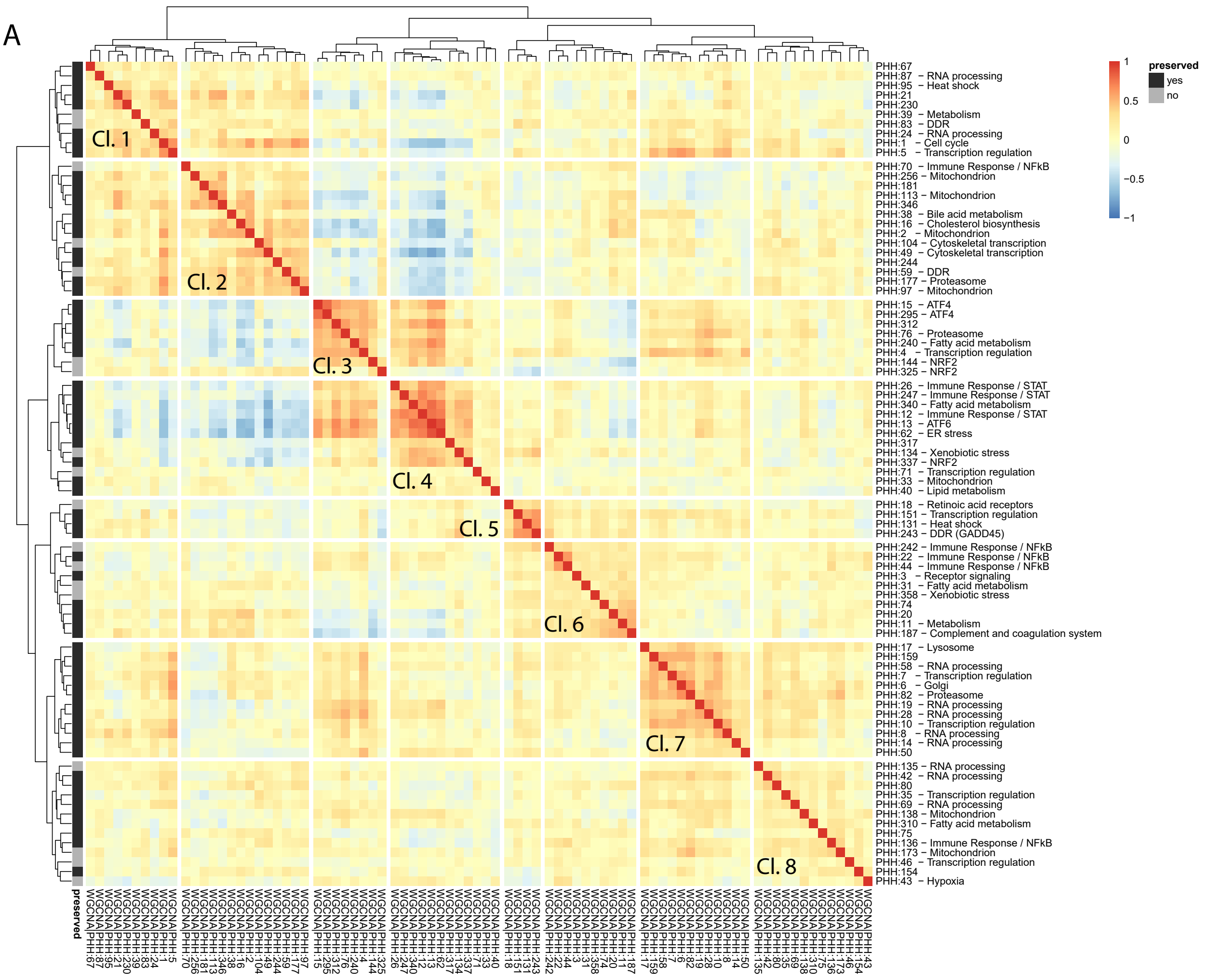


B

**Overlap module clusters**

| | cluster1_s1500 | cluster2_s1500 | cluster3_s1500 | cluster4_s1500 | cluster5_s1500 | cluster6_s1500 | cluster7_s1500 | cluster8_s1500 |
|---|---|---|---|---|---|---|---|---|
| cluster1 TG: Immune Response | 1 (0) | 0.811 (1) | 1 (0) | 0.105 (3) | 1 (0) | 0.064 (3) | 1 (0) | 0.403 (2) |
| cluster2 TG: TF | 0.726 (1) | 1 (0) | 0.64 (1) | 0.793 (1) | 0.392 (1) | 1 (0) | 0.004 (5) | 0.82 (1) |
| cluster3 TG: Stress (ATF4) | 1 (0) | 0.784 (2) | 0.035 (4) | 0.928 (1) | 0.019 (3) | 0.885 (1) | 0.928 (1) | 0.443 (3) |
| cluster4 TG: Stress ATF6, Nrf2) | 1 (0) | 0.873 (1) | 0.266 (2) | 0.041 (4) | 1 (0) | 0.37 (2) | 0.825 (1) | 0.851 (1) |
| cluster5 TG: Mitochondria | 1 (0) | 0.114 (3) | 1 (0) | 0.711 (1) | 1 (0) | 1 (0) | 1 (0) | 0.094 (3) |
| cluster6 TG: RNA processing | 0.005 (4) | 0.77 (1) | 1 (0) | 1 (0) | 1 (0) | 1 (0) | 0.304 (2) | 0.742 (1) |
| cluster7 TG: Mix | 0.046 (3) | 0.77 (1) | 1 (0) | 1 (0) | 1 (0) | 1 (0) | 0.076 (3) | 1 (0) |
| cluster8 TG: Stress (DDR, fatty acid | 0.625 (2) | 0.101 (5) | 0.838 (1) | 0.733 (2) | 1 (0) | 0.1 (4) | 1 (0) | 0.777 (2) |
